# Quantifying irreversibility of ecological systems

**DOI:** 10.1101/2024.08.04.606544

**Authors:** Junang Li, Stephan B. Munch, Tzer Han Tan, Chuliang Song

## Abstract

Irreversibility—the asymmetry of population dynamics when played forward versus backward in time—is a fundamental property of ecological dynamics. Despite its early recognition in ecology, irreversibility has remained a high-level and unquantifiable concept. Here, we introduce a quantitative framework rooted in non-equilibrium statistical physics to measure irreversibility in general ecological systems. Through theoretical analyses, we demonstrate that irreversibility quantifies the degree to which a system is out of equilibrium, a property not captured by traditional ecological metrics. We validate this prediction empirically across diverse ecological systems structured by different forces, such as rapid evolution, nutrient availability, and temperature. In sum, our study provides a rigorous formalism for quantifying irreversibility in ecological systems, with the potential to integrate dynamical, energetic, and informational perspectives in ecology.

## Introduction

Time is the relentless sculptor of life on Earth. A defining characteristic of living systems is their asymmetry in time: the trajectory of life differs significantly when played forward versus backward. For example, Dollo’s law of irreversibility proposes that complex traits lost in evolution are rarely regained [2]. This fundamental asymmetry permeates all life forms, as their diversity, distribution, and functioning are all shaped by non-random and directional ecological and evolutionary processes. While irreversibility manifests at all scales of biological organization [3, 4, 5, 6], we focus here on population dynamics of ecological communities.

The concept of irreversibility is central to physics, and a brief exploration of the physicist’s perspective is instructive. Despite our intuitive experience of the arrow of time, most fundamental physical laws—including Newton’s laws of motion, Maxwell’s equations, and the Schrödinger equation—are time-symmetric: If these equations allow a certain series of events to happen, they equally allow the same events to happen in reverse [7]. This apparent contradiction is resolved by the second law of thermodynamics, which introduces the concept of entropy to explain the statistical likelihood of certain events occurring in one direction over another. To illustrate, consider a balloon filled with high-pressure gas inside a box. Over time, the gas escapes the balloon and fills the box. While the fundamental laws of physics allow for the reverse scenario—the gas spontaneously compressing back into the balloon—the statistical probability of this occurring is incredibly low. Thus, the arrow of time emerges from the statistical differences in the forward versus reverse temporal dynamics [8].

Mirroring this idea, we explore irreversibility in population dynamics using two standard theoretical models as examples of the extremes. Logistic growth model posits deterministic growth towards a carrying capacity. Here, the reversed trajectory is statistically distinct, as it lacks the observed population increase, rendering it *completely irreversible* (Figure 1A1). In contrast, neutral theory posits that species abundances follow uncorrelated random walks. In this model, the forward and reversed trajectories are statistically identical, rending it *completely reversible* (Figure 1A2).

**Figure 1:**
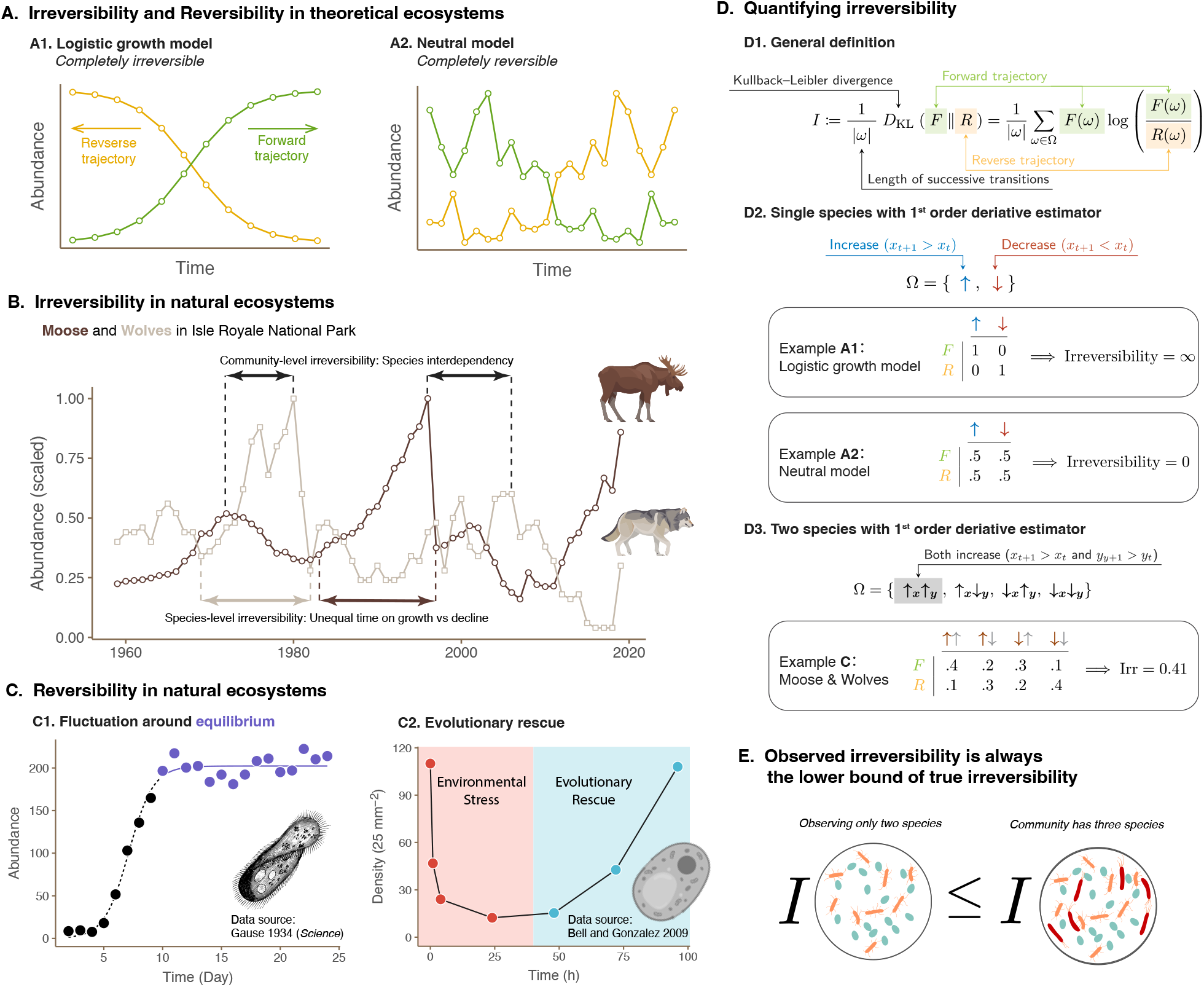
Illustrations of the concept of irreversibility. Panel (**A**) use theoretical models to illustrate extreme cases: (**A1**) the logistic growth model exhibits complete irreversibility, with the forward (green) and reverse (yellow) trajectories being distinct, while (**A2**) the neutral model shows complete reversibility, with statistically identical forward and reverse trajectories. Panel (**B**) shows sources of irreversibility in natural ecosystems, using moose and wolves in Isle Royale National Park as an example [9]. Irreversibility emerges at both the species level, with unequal periods of growth and decline for moose and wolves, and at the community level, with species interdependency leading to peaks in prey abundance preceding those of predators. Panel (**C**) shows sources of reversibility in natural ecosystems: (**C1**) fluctuations around equilibrium [44] and (**C2**) evolutionary rescue in the face of environmental stress [45]. Panel (**D**) illustrates how to quantify irreversibility: (**D1**) The general definition of irreversibility as the Kullback-Leibler divergence between the probability distributions of the forward and reverse trajectories. (**D2**) For a single species, irreversibility is estimated using a first-order derivative estimator based on abundance increases and decreases. Examples illustrate complete irreversibility in the logistic growth model and complete reversibility in the neutral model. (**D3**) For two species, irreversibility is estimated using the joint probabilities of abundance changes across both species, as demonstrated in the wolf and moose system of Isle Royale National Park. Panel (**E**) shows a desirable property of estimating irreversibility from incomplete samplings.

Natural communities, however, occupy the spectrum between these extremes. For example, wolf and moose populations in Isle Royale National Park reveal intricate patterns of irreversibility [9].

At the species level, irreversibility emerges from the unequal time spent on growth and decline, characterized by slow, prolonged growth followed by a rapid, sudden decline (Figure 1B). Similar patterns are observed across ecosystems, such as algal blooms [10], insect outbreaks [11], and coral reef bleaching [12]. At the community level, irreversibility emerges from the interdependency of growth and decline across species. Predation theory predicts that prey populations peak before their predators, a pattern observed in the wolf-moose system that creates community-level irreversibility (Figure 1B).

In addition to these processes promoting irreversibility, ecological processes of natural communities can also reduce irreversibility. For example, after a population reaches the carrying capacity, the subsequent fluctuations around equilibrium are reversible (Figure 1C1). For another example, when a population faces detrimental environments and experiences initial decline, the inherent genetic diversity of a population may enable evolutionary rescue and lead to population growth, exhibiting reduced irreversibility (Figure 1C2).

It is not a new idea that irreversibility is fundamental to ecological systems. Alfred J. Lotka, a founder of modern population biology, proposed characterizing ecological dynamics in terms of irreversibility [13]. Yet, this perspective languished in obscurity, with Lotka’s seminal paper cited a mere five times since its debut a century ago [1]. This neglect partly stems from the challenge of quantifying irreversibility, leaving it a qualitative and vaguely defined concept in ecology.

Recent advancements in non-equilibrium statistical physics offer a canonical description of irreversibility [14, 15]. Leveraging these advances, unavailable in Lotka’s era, we introduce a rigorous, quantitative measure of irreversibility, revitalizing Lotka’s perspective. Our measure of irreversibility is non-parametric, robust to process and measurement noise, and applicable even with unmeasured species. By applying this new approach, we explore how irreversibility captures the dynamics and complexity of ecological systems.

## Quantifying irreversibility

Drawing on the established concept of irreversibility in non-equilibrium statistical physics [16], we introduce a novel metric for quantifying *ecological irreversibility* (*I*). This measure is defined as the Kullback-Leibler (KL) divergence between the probability distributions of observing abundance transitions in a forward trajectory (ℱ) and its time-reversed counterpart (ℛ) (Figure 1D):

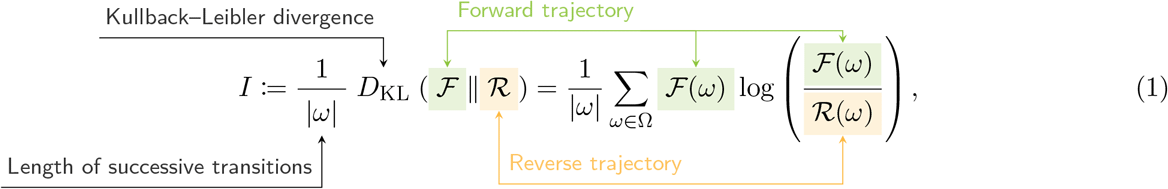

where *ω* represents abundance transitions, Ω is the set of all possible transitions, and |*ω*| denotes the length of successive transitions. A higher *I* indicates greater distinction between forward and reverse trajectories, implying higher irreversibility.

To compute irreversibility, we need to specify the set of abundances transitions (Ω). For a single species, we focus on a coarse-grained scale by considering abundance increases and decreases at each time step (meaning |*ω*| = 1):

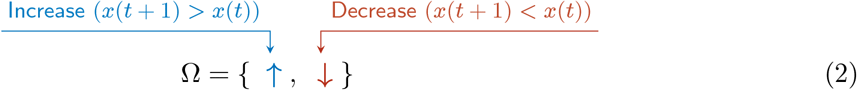

To illustrate its usage, we revisit the two ecological models mentioned earlier (Figure 1A). For logistic growth (Figure 1D2), there is abundance increase in the forward trajectory but not in the reversed trajectory (ℱ (↑)≠0 while ℛ (↑) = 0), implying complete irreversibility (*I* = ∞). Conversely, for neutral theory (Figure 1D2), the probability of an increase or decrease in abundance is equal (ℱ (↑) = ℛ (↑) and ℱ (↓) = ℛ (↓)), implying complete reversibility (*I* = 0).

This approach is readily extendable to multiple species. For a two-species system, we consider all possible combinations of increases and decreases across both species (Ω = {↑_1_↑_2_, ↑_1_↓_2_, ↓_1_↑_2_, ↓_1_↓_2_}). For instance, in the wolf-moose example, irreversibility is estimated at approximately 0.41 by calculating the probabilities of each state in both the forward and reverse trajectories (Figure 1D3).

In the most general case, the successive transition *ω* can be generalized to an event sequence from time 0 to the observation time *T*_obs_ as 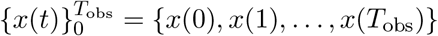. In the physical context, as the observation time *T*_obs_ approaches infinity, the irreversibility metric (Eqn. 1) quantifies the entropy production rate in a nonequilibrium steady state [17]. This most general definition is difficult to estimate directly because it often demands millions of data points [3]. However, ecological time series are typically short and noisy, making the precise estimation impractical. Thus, we prioritize our coarse-grained measure of irreversibility in ecological dynamics.

Despite the simplicity of our approach, using only binary transitions, it still captures the essential patterns of irreversibility, akin to a compressed audio file still conveys the essence of a melody (Appendix C). This is due to the mathematical properties of KL divergence, which enable the construction of various bounds through appropriate coarse-graining schemes. For example, this coarse-grained measure consistently provides a *conservative* (lower bound) estimate of irreversibility compared to measures that incorporate more information (see proof in Appendix A). This property is crucial in ecology data analysis, where incomplete sampling of species is common [18]. While incomplete sampling can introduce unpredictable bias into various time series estimates, such as the Lyapunov exponent, our measure ensures that the observed irreversibility never exceeds the true value for the entire system.

### Irreversibility measures disequilibrium

What insights can our measure of ecological irreversibility (Eqn. 1) provide that other measures cannot? Unlike other measures, irreversibility is not a measure of chaos or stability. A chaotic system, as we demonstrate by theoretical models [19] and empirical analyses of the Global Population Dynamics Database [20], can exhibit varying levels of ecological irreversibility (Appendix D). Similarly, systems with different rates of return to equilibrium can display identical ecological irreversibility (Appendix E).

Instead, ecological irreversibility quantifies the *extent of disequilibrium* within a system, with higher values indicating greater distance from ecological equilibrium. To establish the intuition, we revisit previous empirical examples. Processes that *increase* irreversibility tend to drive directional changes, such as shifting to a new equilibrium (Figure 1A1) or exhibiting unequal periods of growth and decline around an existing equilibrium (Figure 1B). Conversely, processes that *decrease* irreversibility tend to push the system towards equilibrium, such as stochasticity around equilibrium (Figure 1C1) or evolutionary rescue (Figure 1C2). To our knowledge, no other methods can readily extract this information from ecological time series, especially when the precise location of equilibrium is often unknown in empirical data.

Fundamentally, the underlying causal pathway is the inherent asymmetry of ecological processes. When a system is at equilibrium, often referred to as the “balance of nature” [21], opposing forces are precisely balanced, resulting in time-symmetrical and reversible dynamics. However, any deviation from this equilibrium state disrupts this balance, introducing asymmetry into the system. This asymmetry is amplified by the nonlinear nature of ecological interactions, becoming more pronounced as the system moves further from equilibrium. This fundamental asymmetry manifests as measurable irreversibility in the system’s dynamics, thus quantifying the extent of disequilibrium. A more detailed discussion on formalizing the asymmetry in terms of nonreciprocity can be found in Appendix L.

To formalize and demonstrate this conceptual link, we examine three distinct types of ecological systems: 2-species predation dynamics, many-species dynamics with random structure, and many-species dynamics with metabolic structure. Through these examples, we will demonstrate the consistent ability of irreversibility to reflect disequilibrium and reveal insights hidden within ecological data. This approach offers a powerful new tool for understanding the complex dynamics of ecological systems and predicting their responses to perturbations.

#### Predation dynamics

The Lotka-Volterra predation model, a cornerstone in ecological modeling, describes the dynamics between a prey (*N*) and a predator (*P*):

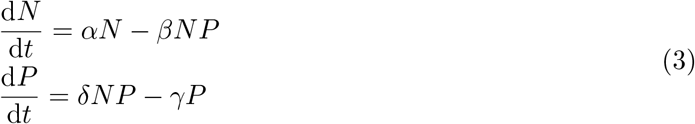

This model produces a persistent cycle (Figure 2A). Within the same system, initial conditions further from equilibrium lead to larger fluctuations, which, as visually observed, result in greater asymmetry in forward and reverse probabilities (e.g., ℱ (**↑**_*N*_ **↑**_*P*_) = ℛ (**↓**_*N*_ **↓**_*P*_) *≠*ℛ (**↓**_*N*_ **↓**_*P*_)). We analytically derive that this intuition holds: the further the system is from equilibrium, the higher its ecological irreversibility (Appendix F.1). This model belongs to a class of models known as conservative ecological dynamics, which allows for the definition of a Hamiltonian (analogous to system energy). The Hamiltonian is directly related to the distance to equilibrium (proof in F.1), aligning our results with the expectation that higher energy drives increased irreversibility. We further confirmed the generality of this relationship by examining other conservative ecological dynamics (Appendix F.2).

**Figure 2:**
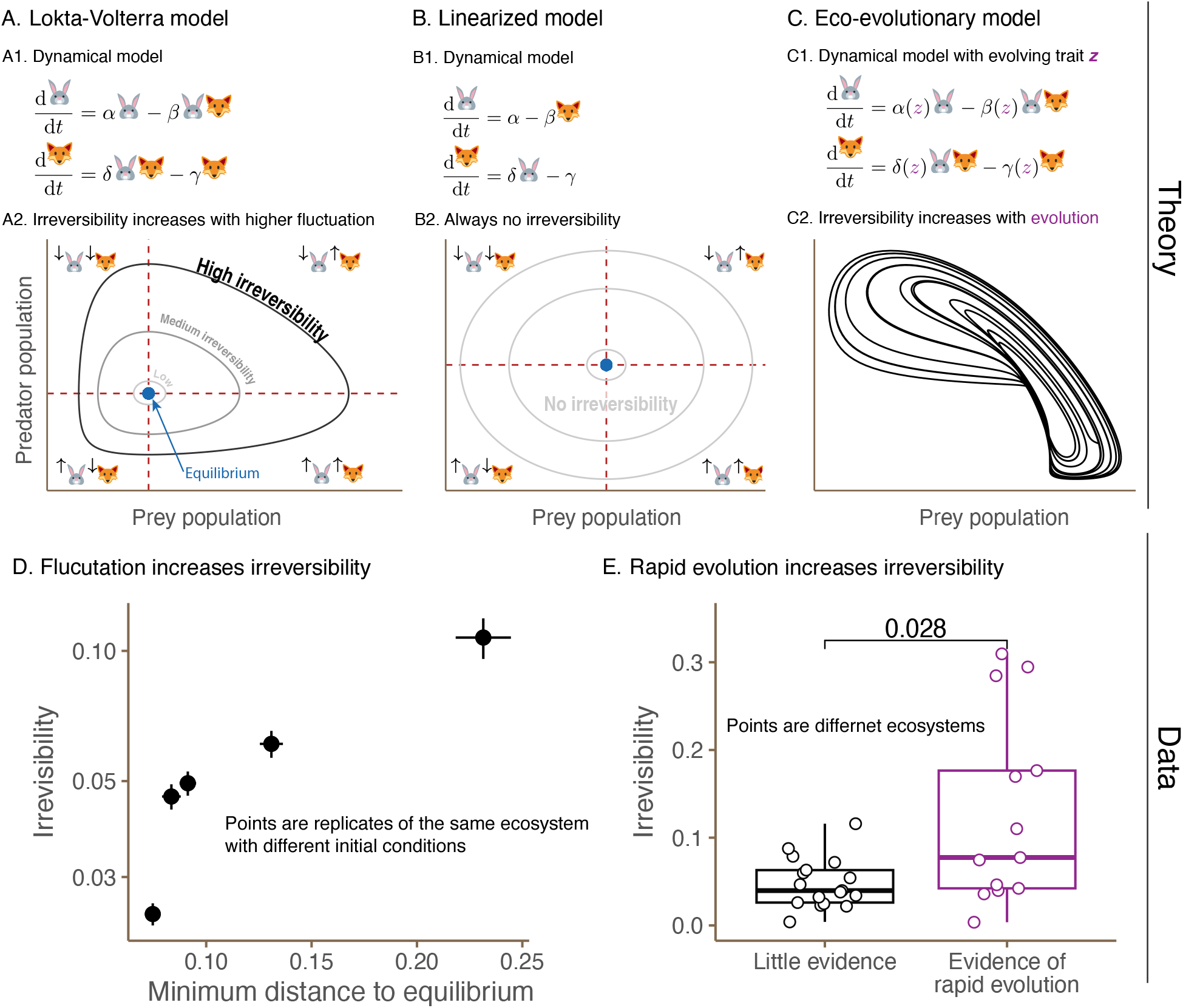
Irreversibility in predation dynamics. Panel (**A**) illustrates the Lotka-Volterra predation model: (**A1**) Dynamical equations describing the interactions between prey and predator populations. (**A2**) Irreversibility increases with higher fluctuations, as the system moves further away from equilibrium (indicated by the blue dot). Panel (**B**) illustrates the linearized version of the Lotka-Volterra model: (**B1**) Dynamical equations of the linearized model. (**B2**) The linearized model always exhibits zero irreversibility under our measure, regardless of the magnitude of fluctuations. Panel (**C**) illustrates the eco-evolutionary predation model: (**C1**) Dynamical equations incorporating an evolving trait *z*, such as body size or defense level. (**C2**) Irreversibility increases with the inclusion of rapid evolution, as the system is driven away from its initial equilibrium. Panel (**D**) tests contrasting predictions of Lotka-Volterra dynamics and its linearized version. Empirical data from a predator-prey system (*Brachionus calyciflorus* and *Monoraphidium minutum*) [24] confirms the theoretical prediction that irreversibility increases with the minimum distance to equilibrium. Points represent four replicates of the same ecosystem with different initial conditions, and the error bars correspond to two standard errors. Panel (**E**) tests whether rapid evolution increases irreversibility. Analysis of multiple empirical datasets reveals that systems with rapid evolution (purple) generally exhibit higher irreversibility compared to those without rapid evolution (black). Points represent different ecosystems, and the distribution of irreversibility values is shown for each category.

Importantly, this relationship between distance from equilibrium and our irreversibility estimator is a characteristic of nonlinear ecological dynamics. The linearized version of the Lotka-Volterra model, initially considered by Lotka [22] and Volterra [23], also produces a persistent cycle (Figure 2B). However, this cycle is always symmetric under our coarse-grainning, meaning that the forward and reverse probabilities of state transitions are always identical (e.g., ℱ(**↑**_*N*_ **↑**_*P*_) = ℛ (**↑**_*N*_ **↑**_*P*_)).

Therefore, under linear dynamics, ecological irreversibility remains zero regardless of fluctuation amplitude (proof in Appendix F.3). Note this property is specific to our coarse-graining approach; the underlying persistent cycle exhibits a directional trend, suggesting irreversibility within this linear system.

To empirically test whether ecological irreversibility increases with the distance to equilibrium, we use long-term experimental data of a planktonic predator–prey system [24]. With five replicates under constant environmental conditions, the data confirmed that irreversibility increases with the distance to equilibrium (Figure 2D; details in Appendix G.1).

Next, we consider predation dynamics with rapid evolution. Trait evolution [25] typically introduces novel dynamics beyond perpetual cycles (Figure 2C). This generally promotes irreversibility, as it drives the system away from its initial equilibrium to some evolving new equilibrium. To empirically test this prediction, we analyzed 18 datasets without rapid evolution [24, 26], and 13 datasets with rapid evolution [26]. Aligning with theoretical prediction, rapid evolution generally increases system’s irreversibility (Figure 2E; details in Appendix G.2).

#### Multispecies dynamics with unstructured interactions

To understand the dynamics of species-rich systems, a canonical approach is to assume that inter-specific interaction strengths are drawn from a random distribution. This assumption is ecologically justified when species interactions emerge from a high-dimensional space of ecological traits [27]. In this line, we consider the stochastic Lotka-Volterra model with dispersal for *S* species:

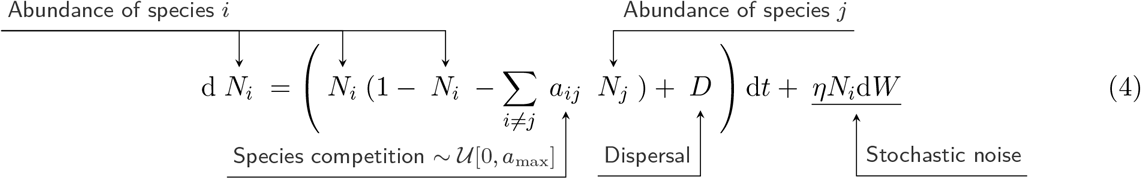

where *N*_*i*_ represents the abundance of species *i, a*_*ij*_ is the competition coefficient between species *i* and *j* (drawn from some distribution), *D* is the dispersal rate, and *η* is the noise level and d*W* is a standard Wiener process.

Ecological theory predicts distinct phases characterized by either low fluctuations driven by stochastic noise (Figure 3A3) or high fluctuations driven by species interactions (Figure 3A2) [28]. These phases are universal, independent of specific settings such as distribution type and functional response [29]. Importantly, the fluctuation phases are determined by species richness and average competition strength, with higher species richness or stronger competition leading to the high-fluctuation phase and vice versa.

**Figure 3:**
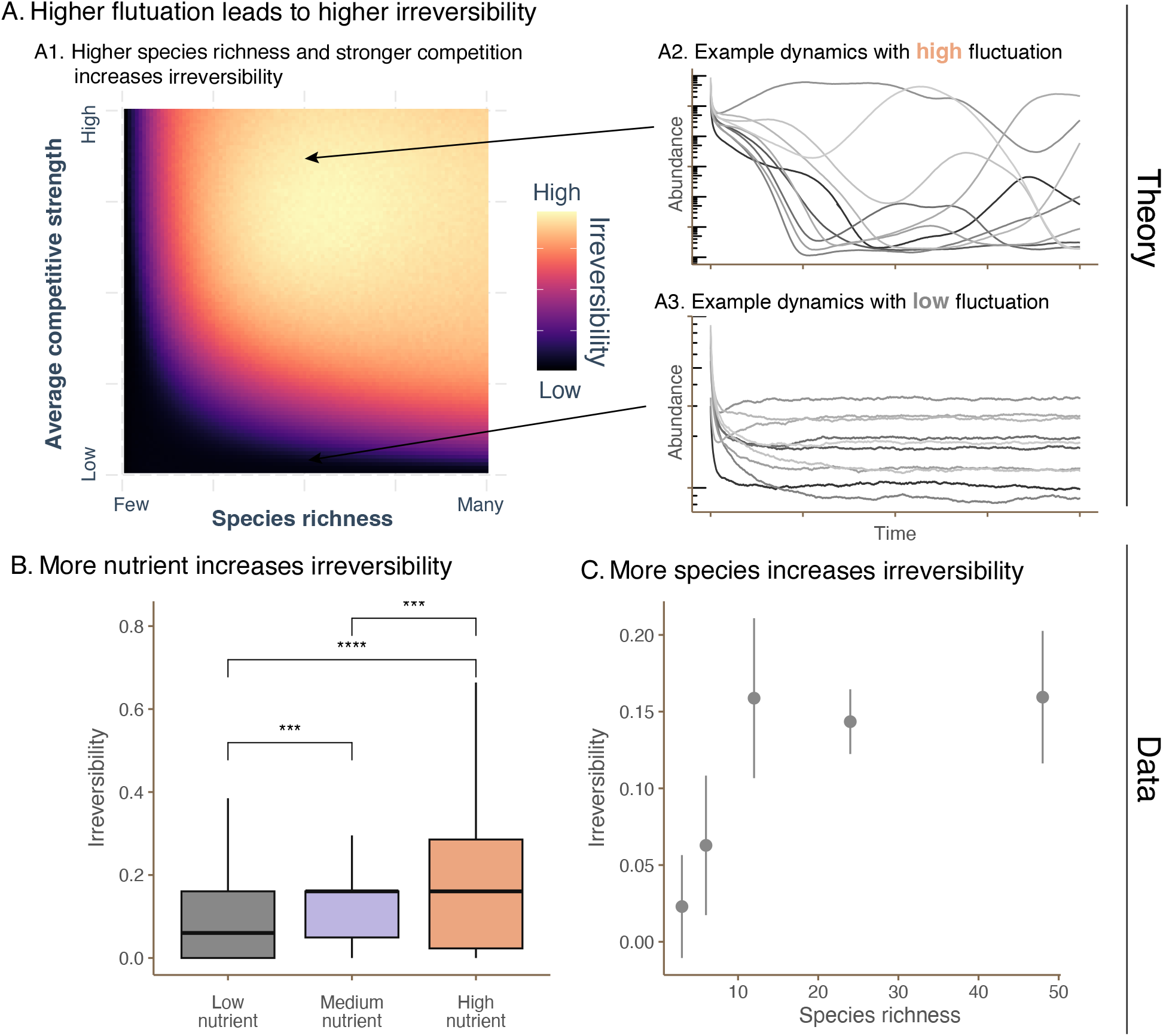
Irreversibility in unstructured multispecies systems. Panel (**A**) illustrates how higher fluctuations lead to higher irreversibility in a stochastic Lotka-Volterra model with dispersal (Eqn. 4). (**A1**) Irreversibility increases with higher species richness and stronger competition, as these factors drive the system into the high fluctuation phase. (**A2**) Example dynamics in the high fluctuation phase exhibit large, irregular oscillations. (**A3**) Example dynamics in the low fluctuation phase show small, noise-driven fluctuations. Panel (**B**) tests the prediction that more nutrients— which increases mean competition strength [31]—increase irreversibility in empirical microbiome data [30]. The boxplots show the distribution of irreversibility values for communities grown in low, medium, and high nutrient conditions. The box represents the 50% of the central data, with a line inside that represents the median. Panel (**C**) tests the prediction that more species increase irreversibility and then saturates using the same empirical microbiome data with low nutrient availability. The scatter plot shows irreversibility as a function of species richness (all under low nutrient conditions), with the points showing the median and the error bars showing two standard errors, revealing a positive relationship.

Linking fluctuation phases to irreversibility, we find that the high-fluctuation phase is characterized by high irreversibility, while the low-fluctuation phase exhibits low irreversibility (Figure 3A1; Appendix H). This finding further supports the general association between disequilibrium and irreversibility. We then test our predictions with empirical microbiome data [30] (Appendix I). In microbiome communities, average interaction strength can be manipulated by altering nutrient levels, with more nutrients leading to stronger competition [31]. Consistent with our theoretical expectations, we find that increased nutrient levels generally lead to higher irreversibility (Figure 3B). Similarly, higher species richness also resulted in increased irreversibility, and this effect saturates with high species richness (Figure 3C).

#### Multispecies dynamics with metabolic constraints

The metabolic rate of an organism is a fundamental biological rate that governs many ecological processes. The Metabolic Theory of Ecology links the metabolic rate (*Q*) of an organism to its body mass (*M*) and temperature (*T*) as follows (Figure 4A) [32]:

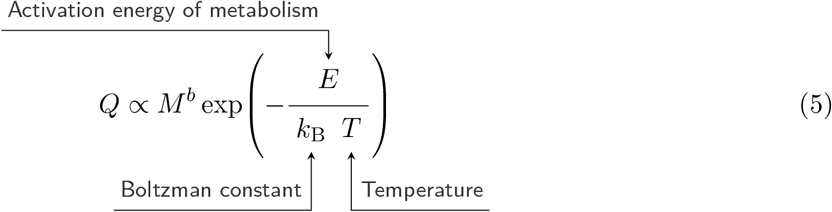

where *E* is the activation energy of metabolism, *k*_B_ is the Boltzmann constant, and *b* is the allometric scaling exponent (∼ 0.75).

**Figure 4:**
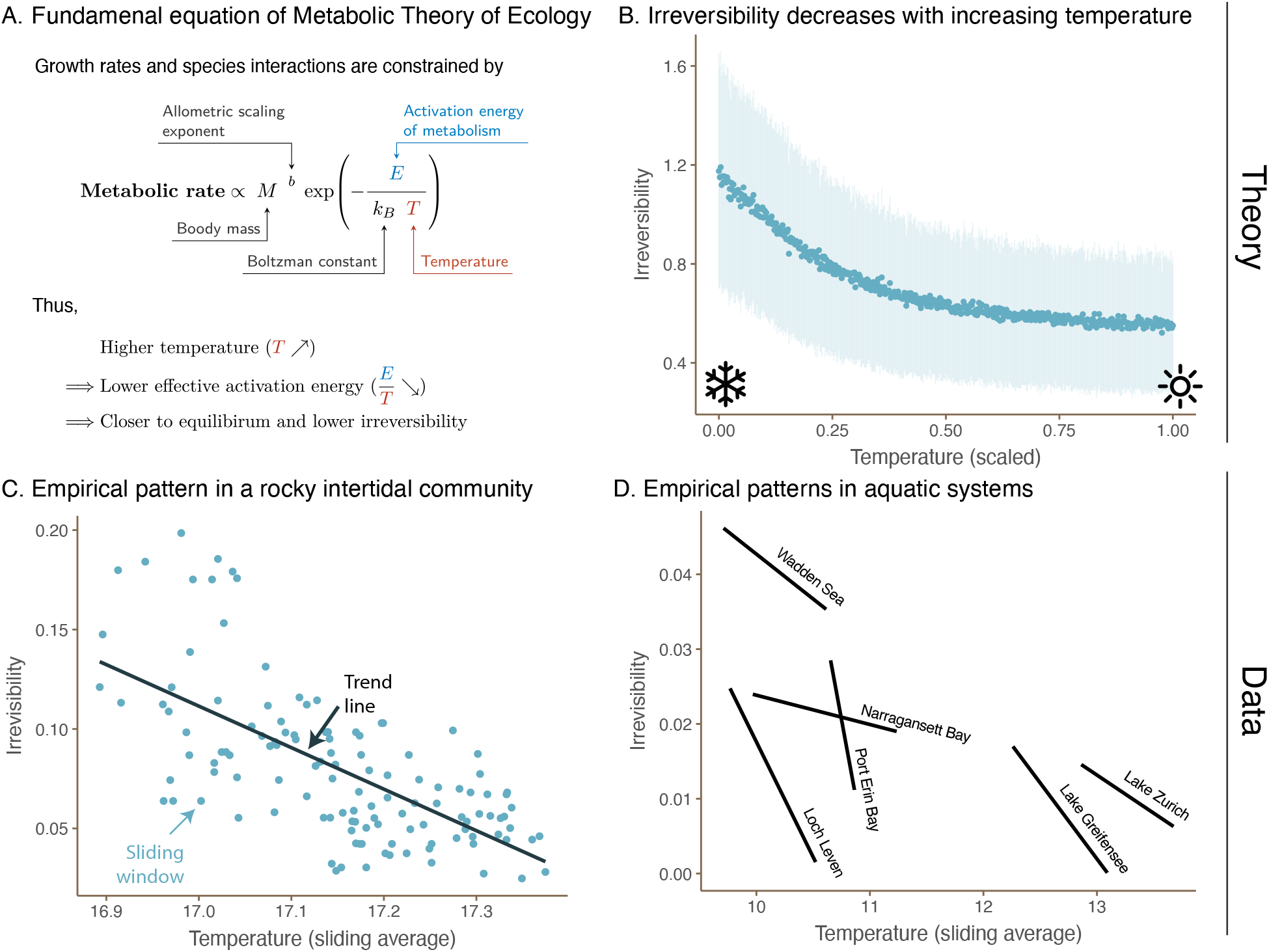
Irreversibility in metabolically-constrained multispecies systems. Panel (**A**) illustrates the Metabolic Theory of Ecology, which links an organism’s metabolic rate to its body mass and temperature (Eqn. 5). Higher temperature effectively lowers the activation energy (*E/T*), bringing the system closer to equilibrium and reducing irreversibility. Panel (**B**) confirms this prediction using simulations of a metabolically-constrained Lotka-Volterra model. The scatter plot shows, with all other parameters fixed, irreversibility decreasing with increasing temperature, with the points representing the median and the error bars showing the 95% confidence interval. Panel (**C**) tests this prediction using empirical data from a rocky intertidal community in Goat Island Bay, New Zealand [34]. Using a sliding window analysis, the scatter plot shows irreversibility decreasing with increasing temperature (sliding average), with the trend line highlighting the negative relationship. Panel (**D**) further corroborates this finding using five additional aquatic datasets, each comprising three species [35]. For each system, irreversibility decreases with increasing temperature, supporting the generality of the temperature-irreversibility relationship predicted by the Metabolic Theory of Ecology.

Since the fundamental equation of the Metabolic Theory of Ecology (Eqn. 5) follows the Arrhenius formalism, temperature modulates the rate rather than the equilibria. However, this local equilibrium description for a single species is altered by the interactions between multiple species. To study how temperature affects irreversibility, we adopt a multispecies Lotka-Volterra model with metabolic constraints [33] (Appendix K)

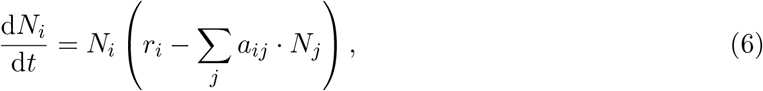

where intrinsic growth rates 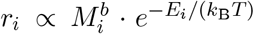, and competition strength 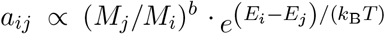. Consequently, increasing the temperature effectively decreases the mean and variance of the competition coefficients. Following previous rationale, temperature increase should result in a lower irreversibility, which is confirmed via simulations (Figure 4B;Appendix K).

We then tested these predictions using empirical data (Appendix J). First, we analyzed a long-term (20+ years) dataset of a rocky intertidal community in Goat Island Bay, New Zealand, which included barnacles, mussels, and algae [34]. A sliding window analysis revealed a decrease in irreversibility with increasing temperature (Figure 4C). To further corroborate this finding, we used six additional datasets, each comprising zooplankton and phytoplankton across various aquatic environments. All datasets confirmed a negative association between temperature and irreversibility [35] (Figure 4D).

## Discussion

Our study introduces a new quantitative framework for measuring irreversibility in ecological systems, a concept long recognized as fundamental but historically challenging to quantify. This measure is firmly rooted in the rich theory of non-equilibrium statistical physics, and is also computationally efficient and applicable to noisy and incomplete observations. Conceptually, our measure of irreversibility has a straightforward interpretation: it serves as a gauge of a system’s distance from ecological equilibrium. The robustness of this interpretation has been rigorously validated through extensive theoretical analyses and emprically tested across diverse ecological systems. Specifically, we have demonstrated that irreversibility increases with factors known to drive systems away from equilibrium, such as rapid evolution, nutrient enrichment, higher biodiversity, and lower temperatures. This new perspective complements current measures and offers a unique lens to uncover the dynamics and complexity of ecological systems.

Irreversibility holds the potential to bridge the long-standing divide between dynamical, energetic, and informational perspectives in ecology. The dynamical perspective, which has long dominated community ecology, models ecological systems as a set of differential equations describing biotic interactions and abiotic factors. In contrast, the informational perspective [36], recently experiencing a resurgence [37, 38], envisions ecosystems as intricate webs of information exchange, with organismal survival hinging on the ability to gather, decipher, and respond to environmental cues. Meanwhile, the energetic perspective [39], central to ecosystem ecology but less prominent in community ecology, emphasizes the acquisition and utilization of energy as the fundamental drivers of organismal interactions and survival. Irreversibility weaves together these seemingly disparate perspectives: it arises through dynamical processes (dynamic), yet functions as an information metric (informational) and, in some cases, serves as the lower bound of energy dissipation (energetic). This unifying potential establishes irreversibility as a central concept for fostering greater integration across ecological subdisciplines.

We acknowledge that our simplified measure cannot fully encapsulate the intricate complexity of irreversibility within ecological systems. For example, the linearized Lotka-Volterra model (Figure 2B) is completely reversible according to our measure, yet one could argue that the persistent cycle with a global trend in the cycle direction signals some level of irreversibility. This discrepancy highlights a key feature of our metric: it aggregates local information (first-order) but not global patterns (infinitely high order; Appendix A). In fact, local and global irreversibility can reflect distinct ecological processes, as irreversibility is inherently scale-dependent [13, 40]. For example, in the Lotka-Volterra dynamics, the global irreversibility (the cycle direction) reflects who is eating whom.

We believe that our work only marks the beginning of a renewed connection between ecology and statistical physics. Ecological theory has predominantly relied on equilibrium-based approaches, rooted in the historical success of merging ecology with *equilibrium* statistical physics [41]. Despite their successes, these approaches increasingly struggle to capture the complexities of ecological systems in a rapidly changing world [42]. Looking forward, recent advances in *non-equilibrium* statistical physics (reviewed in [43]) offers a fertile ground for ecology. By embracing these tools, we may move beyond equilibrium assumptions and gain a deeper understanding of ecological systems far from equilibrium.

## Data and materials availability

All data used in this study are publicly available. Moose and wolf data (Figure 1) are from the Isle Royale Wolf Project (isleroyalewolf.org/data/ data/home.html). Logistic growth data (Figure 1) are from [44] via the gauseR package (CRAN. R-project.org/package=gauseR). Evolutionary rescues data (Figure 1) are from [45] (doi.org/ 10.1111/j.1461-0248.2009.01350.x). Predation dynamics datasets (Figure 2) are from [26] (doi.org/10.1111/ele.12291) and from [24] (figshare.com/articles/dataset/Time_series_of_long-term_experimental_predator-prey_cycles/10045976). Microbiome data (Figure 3) are from [30] (zenodo.org/records/7017202). Rocky intertidal community data (Figure 4) are from [34] (doi.org/10.1073/pnas.1421968112).

