## Supplementary Information for "Quantifying irreversibility of ecological systems"

### Supplementary Material for
#### Quantifying irreversibility of ecological systems

Junang Li[1], Stephan B. Munch[2], Tzer Han Tan[3], and Chuliang Song[4,†]

[1]Center for the Physics of Biological Function,
Princeton University, Princeton, NJ

[2]Department of Applied Mathematics,
University of California, Santa Cruz, CA

[3]Department of Physics,
University of California, San Diego, CA

[4]Department of Ecology and Evolutionary Biology,
University of California, Los Angeles, CA

#### Contents

**A Derivative estimator of irreversibility** **S2**

**B Proof of general properties of irreversibility** **S4**

**C Derivation of irreversibility in neutrality-class model** **S7**

**D Irreversibility does not measure chaos** **S9**

**E Irreversibility does not measure stability** **S14**

**F Theoretical analysis in predation dynamics** **S16**

**G Empirical analysis in predation dynamics** **S22**

**H Theoretical analysis in multispecies dynamics with unstructured interaction** **S29**

**I Empirical analysis in multispecies dynamics with unstructured interaction** **S31**

**J Theoretical analysis in multispecies dynamics with metabolic constraints** **S34**

**K Empirical analysis in multispecies dynamics with metabolic constraints** **S37**

**L Nonreciprocity and irreversibility** **S41**

**M Table of symbols** **S42**

#### A Derivative estimator of irreversibility

We denote a time series trajectory from time 0 to time $T_{\mathrm{obs}}$ with the notation $\{x(t)\}_0^{T_{\mathrm{obs}}} = \{x(0), x(1), \ldots, x(T_{\mathrm{obs}})\}$. We define the true irreversibility ($I_{\mathrm{true}}$) as the Kullback–Leibler divergence between the forward trajectory probability and its reversed trajectory probability,

$$I_{\mathrm{true}} = \lim_{T_{\mathrm{obs}}\to\infty} \frac{1}{T_{\mathrm{obs}}} D_{\mathrm{KL}}(\mathcal{F}(\{x(t)\}_0^{T_{\mathrm{obs}}})||\mathcal{R}(\{x(t)\}_{T_{\mathrm{obs}}}^{0})) \tag{S1}$$

$$= \lim_{T_{\mathrm{obs}}\to\infty} \frac{1}{T_{\mathrm{obs}}} \sum_{x(0),\ldots,x(T_{\mathrm{obs}})} \mathcal{F}(\{x(t)\}_0^{T_{\mathrm{obs}}}) \log \frac{\mathcal{F}(\{x(t)\}_0^{T_{\mathrm{obs}}})}{\mathcal{R}(\{x(t)\}_{T_{\mathrm{obs}}}^{0})} \tag{S2}$$

$$= \lim_{T_{\mathrm{obs}}\to\infty} \frac{1}{T_{\mathrm{obs}}} \sum_{x(0),\ldots,x(T_{\mathrm{obs}})} \mathcal{F}(\{x(0), \ldots, x(T_{\mathrm{obs}})\}) \log \frac{\mathcal{F}(\{x(0), \ldots, x(T_{\mathrm{obs}})\})}{\mathcal{R}(\{x(T_{\mathrm{obs}}), \ldots, x(0)\})}. \tag{S3}$$

Importantly, the probability of a trajectory is equivalent to the probability of observing its temporal derivatives with the same initial condition:

$$\mathcal{P}(\{x(0), \ldots, x(T_{\mathrm{obs}})\}) = \mathcal{P}(\{x(0), x(1) - x(0), \ldots, x(T_{\mathrm{obs}}) - x(T_{\mathrm{obs}} - 1)\}), \tag{S4}$$

This allows us to reformulate the irreversibility in terms of temporal derivatives, denoted as $\Delta x(t) = x(t) - x(t-1)$

$$I_{\mathrm{true}} = \lim_{T_{\mathrm{obs}}\to\infty} \frac{1}{T_{\mathrm{obs}}} \sum_{x(0),\Delta x(1),\ldots,\Delta x(T_{\mathrm{obs}})} \mathcal{F}(\{x(0), \Delta x(1), \ldots, \Delta x(T_{\mathrm{obs}})\}) \log \frac{\mathcal{F}(\{x(0), \Delta x(1), \ldots, \Delta x(T_{\mathrm{obs}})\})}{\mathcal{R}(\{x(T_{\mathrm{obs}}), -\Delta x(T_{\mathrm{obs}}), \ldots, -\Delta x(1)\})}, \tag{S5}$$

Applying the chain rule of KL divergence, we derive:

$$I_{\mathrm{true}} = \lim_{T_{\mathrm{obs}}\to\infty} \frac{1}{T_{\mathrm{obs}}} \sum_{x(0),\Delta x(1),\ldots,\Delta x(T_{\mathrm{obs}})} \mathcal{F}(\{x(0), \Delta x(1), \ldots, \Delta x(T_{\mathrm{obs}})\}) \log \frac{\mathcal{F}(\{x(0), \Delta x(1), \ldots, \Delta x(T_{\mathrm{obs}})\})}{\mathcal{R}(\{x(T_{\mathrm{obs}}), -\Delta x(T_{\mathrm{obs}}), \ldots, -\Delta x(1)\})} \tag{S6}$$

$$= \lim_{T_{\mathrm{obs}}\to\infty} \frac{1}{T_{\mathrm{obs}}} \sum_{\Delta x(1),\ldots,\Delta x(T_{\mathrm{obs}})} \mathcal{F}(\{\Delta x(t)\}_1^{T_{\mathrm{obs}}}) \log \frac{\mathcal{F}(\{\Delta x(t)\}_1^{T_{\mathrm{obs}}})}{\mathcal{R}(\{-\Delta x(t)\}_{T_{\mathrm{obs}}}^{1})} + \tag{S7}$$

$$\sum_{x(0)|\{\Delta x(t)\}_1^{T_{\mathrm{obs}}}} \mathcal{F}(x(0) \mid \{\Delta x(t)\}_1^{T_{\mathrm{obs}}}) \log \frac{\mathcal{F}(x(0) \mid \{\Delta x(t)\}_1^{T_{\mathrm{obs}}})}{\mathcal{R}(x(T_{\mathrm{obs}}) \mid \{-\Delta x(t)\}_{T_{\mathrm{obs}}}^{1})}] \tag{S8}$$

$$\geq \lim_{T_{\mathrm{obs}}\to\infty} \frac{1}{T_{\mathrm{obs}}} \sum_{\Delta x(1),\ldots,\Delta x(T_{\mathrm{obs}})} \mathcal{F}(\{\Delta x(t)\}_1^{T_{\mathrm{obs}}}) \log \frac{\mathcal{F}(\{\Delta x(t)\}_1^{T_{\mathrm{obs}}})}{\mathcal{R}(\{-\Delta x(t)\}_{T_{\mathrm{obs}}}^{1})}. \tag{S9}$$

We define the right-hand side of Equation S9 as the irreversibility of the derivative trajectory ($I_{\mathrm{dev}}$). This establishes $I_{\mathrm{dev}}$ as a lower bound for the true irreversibility, with equality holding if and only if the initial condition has no influence on the trajectory probability.

To enhance computational tractability, we can invoke the Markov assumption and restrict the probability to $d$-th order derivatives:

$$I_{\mathrm{Markov}}^{d} = \frac{1}{d} \sum_{\Delta x(1),\ldots,\Delta x(d)} \mathcal{F}(\{\Delta x(t)\}_1^{d}) \log \frac{\mathcal{F}(\{\Delta x(t)\}_1^{d})}{\mathcal{R}(\{-\Delta x(t)\}_{d}^{1})} \tag{S10}$$

Clearly, $I_{\text{Markov}}$ is a lower bound of $I_{\text{dev}}$. However, $I_{\text{Markov}}$ do not typically converge with limited data (less than 1,000 time steps). Therefore, we introduce a coarse-graining procedure on the derivatives:

$$I_{\text{sign}}^{d} = \frac{1}{d} \sum_{\text{sign}(\Delta x(1)),\ldots,\text{sign}(\Delta x(d))} \mathcal{F}(\{\text{sign}(\Delta x_1), \ldots, \text{sign}(\Delta x_d)\})) \tag{S11}$$

$$\log \frac{\mathcal{F}(\{\text{sign}(\Delta x(1)), \ldots, \text{sign}(\Delta x(d)\}))}{\mathcal{R}(\{\text{sign}(-\Delta x(d)), \ldots, \text{sign}(-\Delta x(1))\})} \tag{S12}$$

As proved in Appendix B, $I_{\text{sign}}^{d}$ is a lower bound of $I_{\text{Markov}}^{d}$.

In summary, we have established a hierarchy of irreversibility measures with varying levels of coarse-graining:

$$I_{\text{true}} \geq I_{\text{dev}} \geq I_{\text{Markov}}^{d} \geq I_{\text{sign}}^{d}. \tag{S13}$$

In the main text, due to data limitations, we have employed $I_{\text{sign}}^{d}$ to quantify irreversibility in empirical data, as it necessitates the least amount of data for convergence.

### B Proof of general properties of irreversibility

#### B.1 Irreversibility is additive with no species interaction

Suppose we have $S$ species, $\mathbf{x} = (x_1, x_2, \ldots, x_S)^T$, that are not interacting with others. In general, the irreversibility $I$ follows that

$$I = \lim_{T_{\text{obs}} \to \infty} \frac{1}{T_{\text{obs}}} \sum_{\{\mathbf{x}(t)\}_0^{T_{\text{obs}}} \in \mathcal{X}} \mathcal{F}(\{\mathbf{x}(t)\}_0^{T_{\text{obs}}}) \log \left( \frac{\mathcal{F}(\{\mathbf{x}(t)\}_0^{T_{\text{obs}}})}{\mathcal{R}(\{\mathbf{x}(t)\}_{T_{\text{obs}}}^0)} \right) \tag{S14}$$

$$= \lim_{T_{\text{obs}} \to \infty} \frac{1}{T_{\text{obs}}} \sum_{\{\mathbf{x}(t)\}_0^{T_{\text{obs}}} \in \mathcal{X}} \mathcal{F}(\{x_1(t)\}_0^{T_{\text{obs}}}, \ldots, \{x_S(t)\}_0^{T_{\text{obs}}}) \log \left( \frac{\mathcal{F}(\{x_1(t)\}_0^{T_{\text{obs}}}, \ldots, \{x_S(t)\}_0^{T_{\text{obs}}})}{\mathcal{R}(\{x_1(t)\}_{T_{\text{obs}}}^0, \ldots, \{x_S(t)\}_{T_{\text{obs}}}^0)} \right) \tag{S15}$$

$$= \lim_{T_{\text{obs}} \to \infty} \frac{1}{T_{\text{obs}}} \sum_{\{\mathbf{x}(t)\}_0^{T_{\text{obs}}} \in \mathcal{X}} \left( \prod_{j=1}^{S} \mathcal{F}(\{x_j(t)\}_0^{T_{\text{obs}}}) \right) \sum_{i=1}^{S} \log \left( \frac{\mathcal{F}(\{x_i(t)\}_0^{T_{\text{obs}}})}{\mathcal{R}(\{x_i(t)\}_{T_{\text{obs}}}^0)} \right) \tag{S16}$$

$$= \sum_{i=1}^{S} \lim_{T_{\text{obs}} \to \infty} \frac{1}{T_{\text{obs}}} \sum_{\{\mathbf{x}(t)\}_0^{T_{\text{obs}}} \in \mathcal{X}} \left( \prod_{j \neq i} \mathcal{F}(\{x_j(t)\}_0^{T_{\text{obs}}}) \right) \mathcal{F}(\{x_i(t)\}_0^{T_{\text{obs}}}) \log \left( \frac{\mathcal{F}(\{x_i(t)\}_0^{T_{\text{obs}}})}{\mathcal{R}(\{x_i(t)\}_{T_{\text{obs}}}^0)} \right) \tag{S17}$$

$$= \sum_{i=1}^{S} \left( \sum_{\mathbf{x} \in \mathcal{X}} \prod_{j \neq i} \mathcal{F}(\{x_j(t)\}_0^{T_{\text{obs}}}) \right) \left( \lim_{T_{\text{obs}} \to \infty} \frac{1}{T_{\text{obs}}} \sum_{\{x_i\}_0^T \in \mathcal{X}} F(\{x_i(t)\}_0^{T_{\text{obs}}}) \log \left( \frac{\mathcal{F}(\{x_i(t)\}_0^{T_{\text{obs}}})}{\mathcal{R}(\{x_i(t)\}_{T_{\text{obs}}}^0)} \right) \right) \tag{S18}$$

$$= \sum_{i=1}^{S} \left( \lim_{T_{\text{obs}} \to \infty} \frac{1}{T_{\text{obs}}} \sum_{\{x_i(t)\}_0^{T_{\text{obs}}} \in \mathcal{X}} F(\{x_i(t)\}_0^{T_{\text{obs}}}) \log \left( \frac{\mathcal{F}(\{x_i(t)\}_0^{T_{\text{obs}}})}{\mathcal{R}(\{x_i(t)\}_{T_{\text{obs}}}^0)} \right) \right). \tag{S19}$$

We define irreversibility $I_i$ of individual species $i$ as

$$I_i = \lim_{T_{\text{obs}} \to \infty} \frac{1}{T_{\text{obs}}} \sum_{x_i \in \mathcal{X}} \mathcal{F}(\{x_i(t)\}_0^{T_{\text{obs}}}) \log \left( \frac{\mathcal{F}(\{x_i(t)\}_0^{T_{\text{obs}}})}{\mathcal{R}(\{x_i(t)\}_{T_{\text{obs}}}^0)} \right). \tag{S20}$$

It is obvious that irreversibility is additive without species interaction

$$I = \sum_{i=1}^{S} I_i \tag{S21}$$

From now on, we will drop the time series notation for notation simplicity. $x_i$ will be used to represent the entire trajectory $\{x_i(t)\}_0^T$ unless specifically notified.

#### B.2 Irreversibility does not decrease with finer scale

Suppose $\bar{\mathcal{X}}$ is a finer probability space of $\mathcal{X}$, and the irreversibility on this finer scale is $\bar{I}$. Suppose each probability $x_i$ in $\mathcal{X}$ is divided into sub-probability $x_i^k$, where $\sum_k x_i^k = x_i$. Then we have

$$I = \lim_{T_{\text{obs}} \to \infty} \frac{1}{T_{\text{obs}}} \sum_{x_i \in \mathcal{X}} \mathcal{F}(x_i) \log \left( \frac{\mathcal{F}(x_i)}{\mathcal{R}(x_i)} \right) \tag{S22}$$

$$= \lim_{T_{\text{obs}}\to\infty} \frac{1}{T_{\text{obs}}} \sum_{\mathbf{x}\in\mathcal{X}} \left(\sum_k \mathcal{F}(x_i^k)\right) \log\left(\frac{\left(\sum_k \mathcal{F}(x_i^k)\right)}{\left(\sum_k \mathcal{R}(x_i^k)\right)}\right) \tag{S23}$$

$$\leq \lim_{T_{\text{obs}}\to\infty} \frac{1}{T_{\text{obs}}} \sum_{\mathbf{x}\in\mathcal{X}} \sum_k \left(\mathcal{F}(x_i^k) \log\left(\frac{\mathcal{F}(x_i^k)}{\mathcal{R}(x_i^k)}\right)\right) \tag{S24}$$

$$= \lim_{T_{\text{obs}}\to\infty} \frac{1}{T_{\text{obs}}} \sum_{x_i^k\in\bar{\mathcal{X}}} \mathcal{F}(x_i^k) \log\left(\frac{\mathcal{F}(x_i^k)}{\mathcal{R}(x_i^k)}\right) \tag{S25}$$

$$= \bar{I}, \tag{S26}$$

where the inequality stems from the log sum inequality.

##### B.3 Observed irreversibility is always the lower bound of true irreversibility

Suppose we have $S$ species and we denote them as $\mathbf{x}$. Then we have a new species which we denote as $y$. Then

$$I(\mathbf{x}, y) = \lim_{T_{\text{obs}}\to\infty} \frac{1}{T_{\text{obs}}} \iint F(\mathbf{x}, y) \log\left(\frac{F(\mathbf{x}, y)}{R(\mathbf{x}, y)}\right) \mathrm{d}\mathbf{x}\, \mathrm{d}y \tag{S27}$$

$$= \lim_{T_{\text{obs}}\to\infty} \frac{1}{T_{\text{obs}}} \iint F(y \mid \mathbf{x}) F(\mathbf{x}) \log\left(\frac{F(y \mid \mathbf{x}) F(\mathbf{x})}{R(y \mid \mathbf{x}) R(\mathbf{x})}\right) \mathrm{d}\mathbf{x}\, \mathrm{d}y \tag{S28}$$

$$= \underbrace{\lim_{T_{\text{obs}}\to\infty} \frac{1}{T_{\text{obs}}} \iint F(y \mid \mathbf{x}) F(\mathbf{x}) \log\left(\frac{F(y \mid \mathbf{x})}{R(y \mid \mathbf{x})}\right) \mathrm{d}\mathbf{x}\, \mathrm{d}y}_{I_1} \tag{S29}$$

$$+ \underbrace{\lim_{T_{\text{obs}}\to\infty} \frac{1}{T_{\text{obs}}} \iint F(y \mid \mathbf{x}) F(\mathbf{x}) \log\left(\frac{F(\mathbf{x})}{R(\mathbf{x})}\right) \mathrm{d}\mathbf{x}\, \mathrm{d}y}_{I_2} \tag{S30}$$

Focusing on the first term, we have

$$I_1 = \lim_{T_{\text{obs}}\to\infty} \frac{1}{T_{\text{obs}}} \int F(y \mid \mathbf{x}) \log\left(\frac{R(y \mid \mathbf{x})}{F(y \mid \mathbf{x})}\right) \mathrm{d}y \int F(\mathbf{x})\, \mathrm{d}\mathbf{x} \tag{S31}$$

$$= \lim_{T_{\text{obs}}\to\infty} \frac{1}{T_{\text{obs}}} \int D_{\text{KL}}(F(y \mid \mathbf{x}) \| R(y \mid \mathbf{x})) F(\mathbf{x})\, \mathrm{d}\mathbf{x} \tag{S32}$$

$$\tag{S33}$$

Notice that both $D_{\text{KL}}(F(y \mid \mathbf{x}) \| R(y \mid \mathbf{x}))$ and $F(\mathbf{x})$ are positive, the integral is also positive. $I_1 \geq 0$.

Focusing on the second term, we have

$$I_2 = \lim_{T_{\text{obs}}\to\infty} \frac{1}{T_{\text{obs}}} \int \left(\int F(y \mid \mathbf{x})\, \mathrm{d}y\right) F(\mathbf{x}) \log\left(\frac{F(\mathbf{x}}{R(\mathbf{x})}\right) \mathrm{d}\mathbf{x} \tag{S34}$$

$$= \lim_{T_{\text{obs}}\to\infty} \frac{1}{T_{\text{obs}}} \int F(\mathbf{x}) \log\left(\frac{F(\mathbf{x})}{R(\mathbf{x})}\right) \mathrm{d}\mathbf{x} \tag{S35}$$

$$= I(\mathbf{x}) \tag{S36}$$

Thus, we have the general relationship that $I(\mathbf{x}, y) \geq I(\mathbf{x})$.

### C Derivation of irreversibility in neutrality-class model

To illustrate how different irreversibility estimators yield qualitatively similar results, we consider a tractable community model: a system of non-interacting species with biased, discrete stochastic population dynamics. Specifically, the abundance $x_i^t$ of species $i$ at time $t$ evolves according to:

$$x_i(t+1) = x_i(t) + \mathcal{U}[-1, 1+\delta] \tag{S37}$$

where $\delta$ represents the non-neutrality, with higher values indicating greater deviation from neutrality.

We assume that each species $i$ has an independent probability $p_i$ of increasing and $(1-p_i)$ probability of decreasing. In a general state space with embedding dimension $d$, the probability for species $i$ of to experience $k$ increases and $(d-k)$ decreases is

$$\mathcal{F}(x_i) = p_i^k (1-p_i)^{d-k},$$

with frequency $\binom{d}{k}$. Conversely, the probability of the reverse trajectory (i.e., $(n-k)$ increases and $k$ decreases) is

$$\mathcal{R}(x_i) = (1-p_i)^k p_i^{d-k}.$$

The irreversibility $I_i$ for species $i$ is then calculated as:

$$\begin{aligned}
I_i &= \frac{1}{d} \sum_{x_i \in \mathcal{X}} F(x_i) \log\left(\frac{\mathcal{F}(x_i)}{\mathcal{R}(x_i)}\right) && \text{(S38)}\\
&= \frac{1}{d} \sum_k \binom{d}{k} p_i^k (1-p_i)^{d-k} \log\left(\frac{p_i^k (1-p_i)^{d-k}}{(1-p_i)^k p_i^{d-k}}\right) && \text{(S39)}\\
&= \frac{1}{d} \sum_k \binom{d}{k} p_i^k (1-p_i)^{d-k} (2k-d) \log\left(\frac{p_i}{1-p_i}\right) && \text{(S40)}\\
&= \frac{1}{d} \log\left(\frac{p_i}{1-p_i}\right) (2 \sum_k \binom{d}{k} k p_i^k (1-p_i)^{d-k} - d \sum_k \binom{d}{k} k p_i^k (1-p_i)^{d-k}) && \text{(S41)}\\
&= (2p_i - 1) \log\left(\frac{p_i}{1-p_i}\right) && \text{(S42)}
\end{aligned}$$

The last step utilizes the combinatorial identity:

$$np(p+q)^{n-1} = \sum_{k=0}^{n} k \binom{n}{k} p^k q^{n-k}. \tag{S43}$$

Expressing the non-neutrality as $\delta_i = 2p_i - 1$ (ranging from 0 for neutrality to 1 for complete determinism), the total irreversibility $I$ of an ecological system with $S$ species is given by:

$$I = \sum_{i=1}^{S} \delta_i \log\left(\frac{1+\delta_i}{1-\delta_i}\right) \tag{S44}$$

In the special case of uniform neutrality (i.e., $\delta_i = \delta, \forall i$), we have

$$I = S\delta \log\left(\frac{1+\delta}{1-\delta}\right) \tag{S45}$$

Figure S1 illustrates this scenario, showing that irreversibility generally increases with higher species richness and greater non-neutrality.

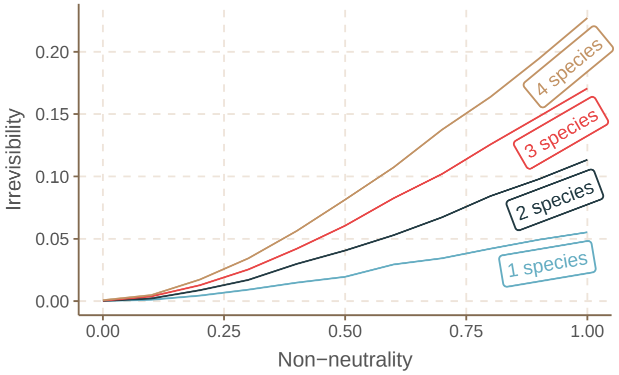


Figure S1: The horizontal axis shows the level of non-neutrality where 0 represents complete neutrality. The vertical axis shows the irreversibility in the system. The four different lines correspond to different species richness in the system. In general, an ecological system becomes more irreversible with higher species richness and higher non-neutrality.

### D Irreversibility does not measure chaos

#### D.1 Heuristic arguments

Two common measures of chaos are Lyapunov exponents and permutation entropy. We argue here why we should expect irreversibility to be fundamentally different from these measures. These heuristic arguments are further supported by simulation studies and empirical analyses, which we detail in the following sections.

We first focus on Lyapunov exponents. The Lyapunov exponent for discrete dynamics $x(t+1) = f(x(t))$ is defined as:

$$\lambda = \lim_{T_{\text{obs}} \to \infty} \frac{1}{T_{\text{obs}}} \sum_{t=0}^{T_{\text{obs}}-1} \ln |f'(x(t))| \tag{S46}$$

This measures the average exponential rate of divergence or convergence of nearby trajectories. Critically, the absolute value of the derivative is used. In contrast, our deriative-absed irreversibility measure explicitly distinguishes between increasing and decreasing trends in the data.

We then focus on permutation entropy. Permutation entropy quantifies the complexity of a time series by considering the relative frequency of different orderings (permutations) of values within an embedding dimension [1]. The metric has used as a measure of chaos in ecological time series [2, 3].

While our measure irreversibility appears to be similar to permutation entropy, they are fundamentally different. For example, consider an embedding dimension of 2. Permutation entropy accounts for six possible orderings of three consecutive points (or, alternately, for two species):

$$123 \rightarrow \nearrow\nearrow; \quad 132\&231 \rightarrow \nearrow\searrow; \quad 213\&312 \rightarrow \searrow\nearrow; \quad 321 \rightarrow \searrow\searrow \tag{S47}$$

Permutation entropy is thus sensitive to a wider range of patterns than irreversibility, which focuses specifically on the balance between increasing and decreasing segments. This distinction becomes even more pronounced with higher embedding dimensions or multiple species.

#### D.2 Simulation analysis with discrete logistic growth model

We consider the classic discrete logistic growth model [4], a canonical example in population dynamics:

$$x(t+1) = \lambda x(t)(1 - x(t)) + \eta x(t) \tag{S48}$$

where $\lambda$ (1 to 4) controls the maximum growth rate, and $\eta$ introduces stochastic noise (e.g., $\mathcal{U}[-0.01, 0.01]$).

This model exhibits a rich range of behaviors, from stable equilibrium to chaotic dynamics, as illustrated by its bifurcation diagram (Figure S2) :

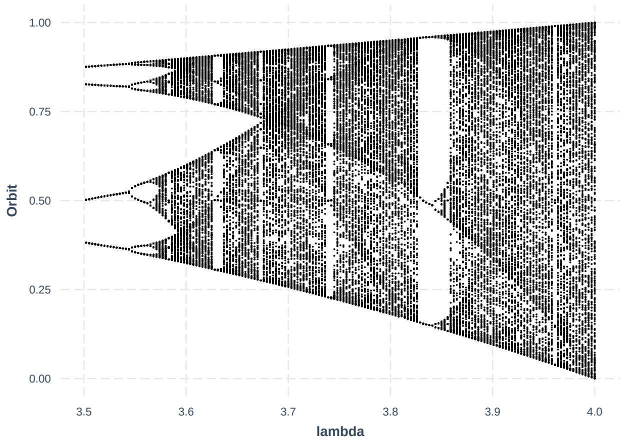


Figure S2: Bifurcation diagram of

Our analysis reveals that irreversibility patterns vary systematically with the growth rate $\lambda$ (Figure S3)

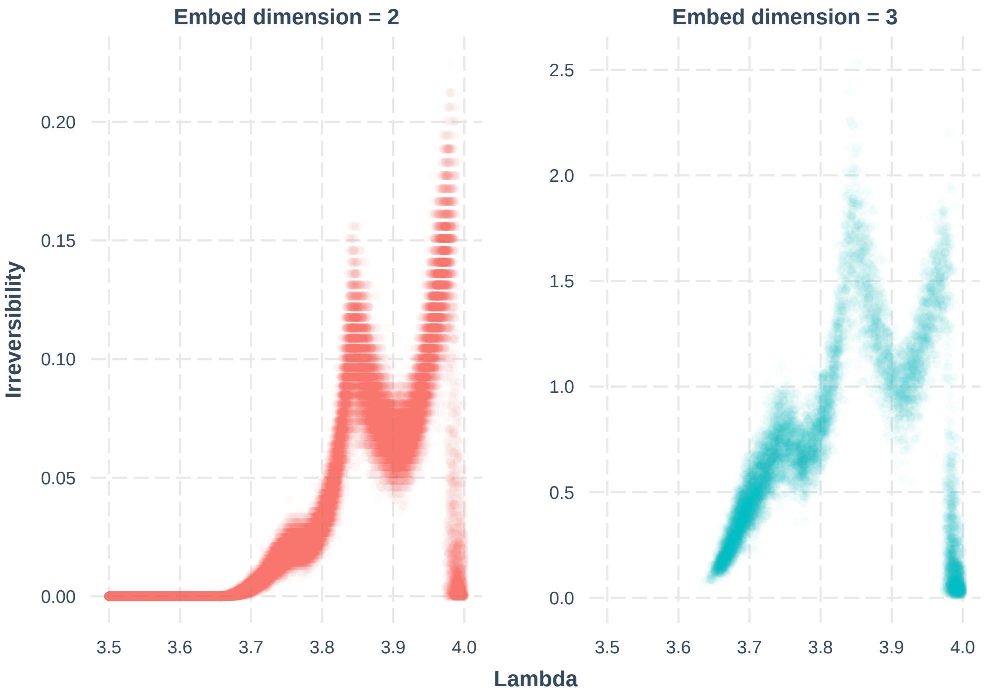


Figure S3: Relationship between irreversibility and lambda in the discrete logistic growth model.

However, these patterns do not align with the Lyapunov exponents (Figure S4). The correlation between irreversibility and Lyapunov exponents is 0.084 with a 95% confidence interval [-0.019, 0.19] when the embedded dimension is 2, and is -0.15 with a 95% confidence interval [-0.024, -0.04] when the embedded dimension is 3.

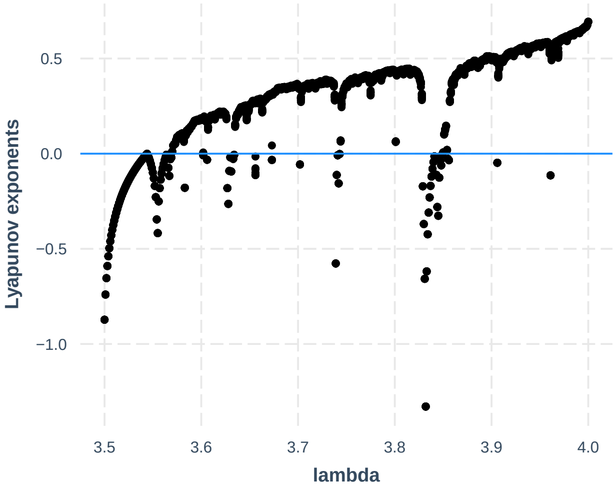


Figure S4: Pattern.

Similarly, the permutation entropy of the system does not correlate with irreversibility (Figure S5). The correlation is 0.09 with a 95% confidence interval [0.08, 0.10] with an embedding dimension of 2, and is 0.30 with [0.29, 0.31].

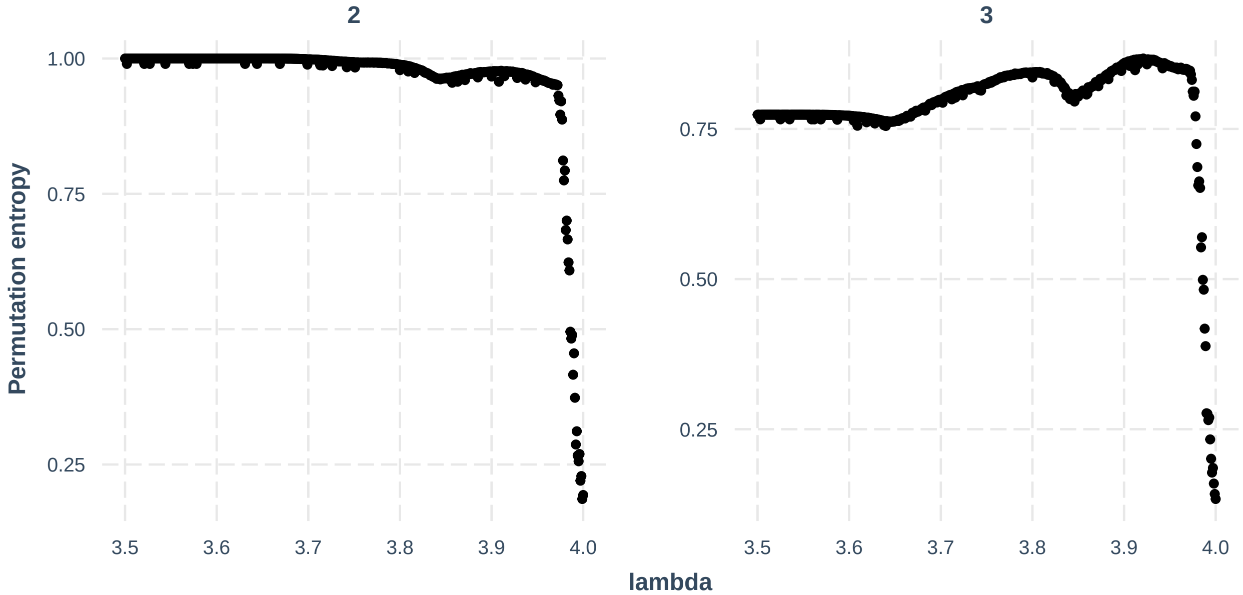


Figure S5: Pattern.

These simulation results further support the conclusion that irreversibility is not a direct measure of chaos.

#### D.3 Empirical Analysis with Global Population Dynamics Database

We further test if our result holds in empirical data.

The Global Population Dynamics Database (GPDD) is currently the largest compilation of time series for single species [5]. As some data is either too short or too noisy for robust time series analysis, we restrict our analysis to a "high-quality" subset [3]. In total, it contains 172 time series spanning 138 different taxa and 57 sampling locations.

We first show its convergence to ensure that time series is long enough to well estimate the values (Figure S6):

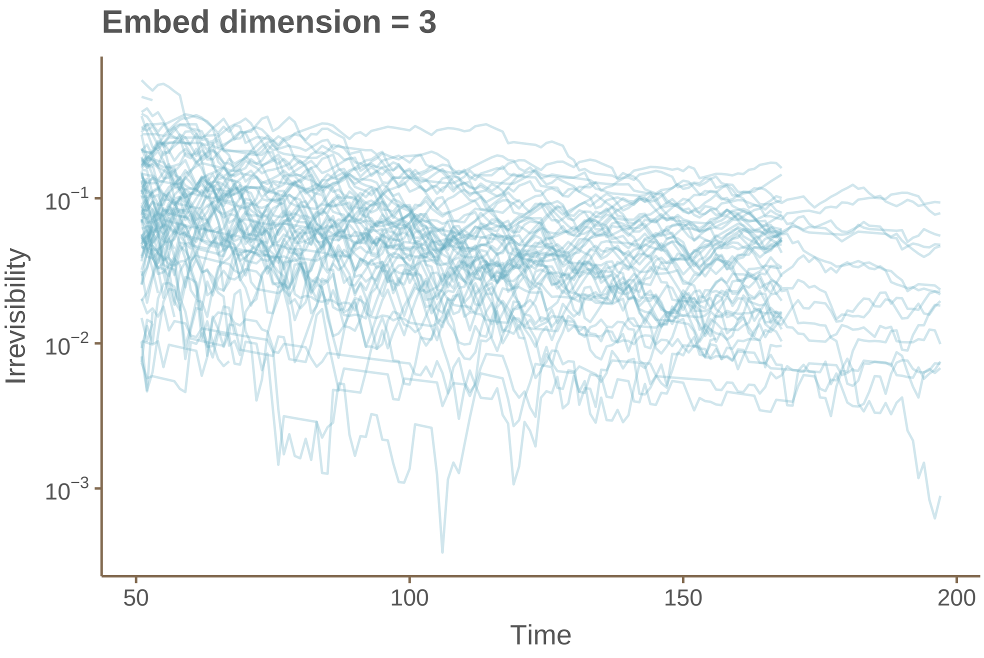


Figure S6: Embed dimension = 3.

Figure S7 shows the association between irreversibility and the Lyapunov exponent in the network. The correlation is insignificant ($-0.068$ with p-value 0.38).

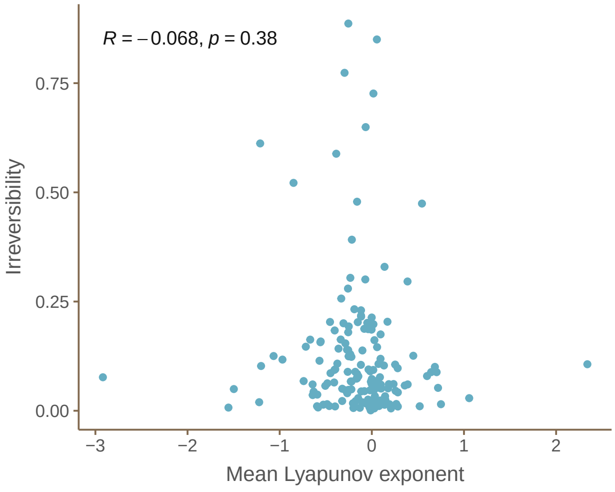


Figure S7: Association between Lyapiunov exponent and irreversibility across datasets in The Global Population Dynamics Database.

Figure S8 shows the association between irreversibility and perturbation in the network. The correlation is also insignificant (−0.085 with p-value 0.27).

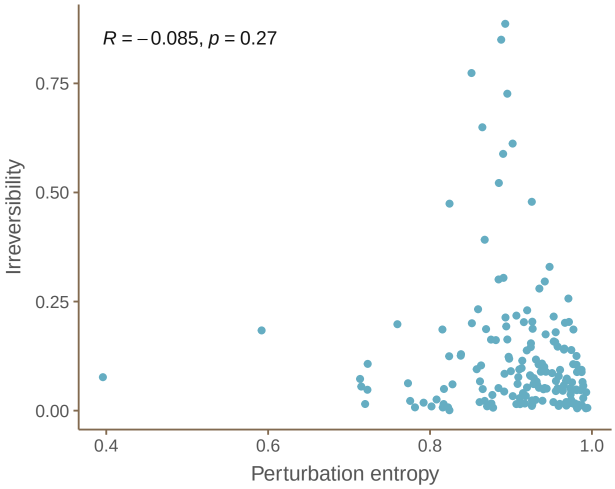


Figure S8: Association between Lyapiunov exponent and irreversibility across datasets in The Global Population Dynamics Database.

#### D.4 A short note on deterministic logistic map

It's worth noting that even the deterministic logistic map at its most chaotic (with $\lambda = 4$) exhibits complete irreversibility when considering 2nd-order embedding. This is because the dynamics never produce two consecutive declines, as can be proven mathematically.

This is because that the dynamics would never have two consecutive declines. That is, there does not exist a time step $t$ such that $x(t+1) < x(t)$ and $x(t+2) < x(t+1)$. The proof is straightforward. If $x(t+1) < x(t)$, then this is equivalent to $x(t) > 3/4$. Similarly, $x(t+2) < x(t+1)$ implies that $x(t+1) > 3/4$. However, if $x(t) > 3/4$, then $x(t+1) < 4 \times 3/4 \times (1 - 3/4) = 3/4$, which causes a contradiction.

### E Irreversibility does not measure stability

Here, we consider stability as the return rate, which is the largest real part of the eigenvalues of the Jacobian matrix. Specifically, for general ecological dynamics,

$$\frac{\mathrm{d}\mathbf{N}}{\mathrm{d}t} = f(\mathbf{N}) \tag{S49}$$

where $\mathbf{N}$ is the vector representing species abundance and $f$ is some arbitrary population dynamics. The Jacobian $\mathbf{J}$ is

$$J = \frac{\partial \frac{\mathrm{d}\mathbf{N}}{\mathrm{d}t}}{\partial \mathbf{N}} \tag{S50}$$

which we can evaluate at equilibrium or out of equilibrium [6, 7].

#### E.1 Simulation with single species

We consider the canonical logistic growth model,

$$\frac{\mathrm{d}N}{\mathrm{d}t} = rN(\frac{K-N}{K}) \tag{S51}$$

The largest eigenvalue at equilibrium is

$$\lambda = -r \tag{S52}$$

Without loss of generality, we choose the carrying capacity $K = 1$. We ran a stochastic simulation at equilibrium that varies $r$ (changing resilience). Figure S9 shows a null association between resilience and irreversibility.

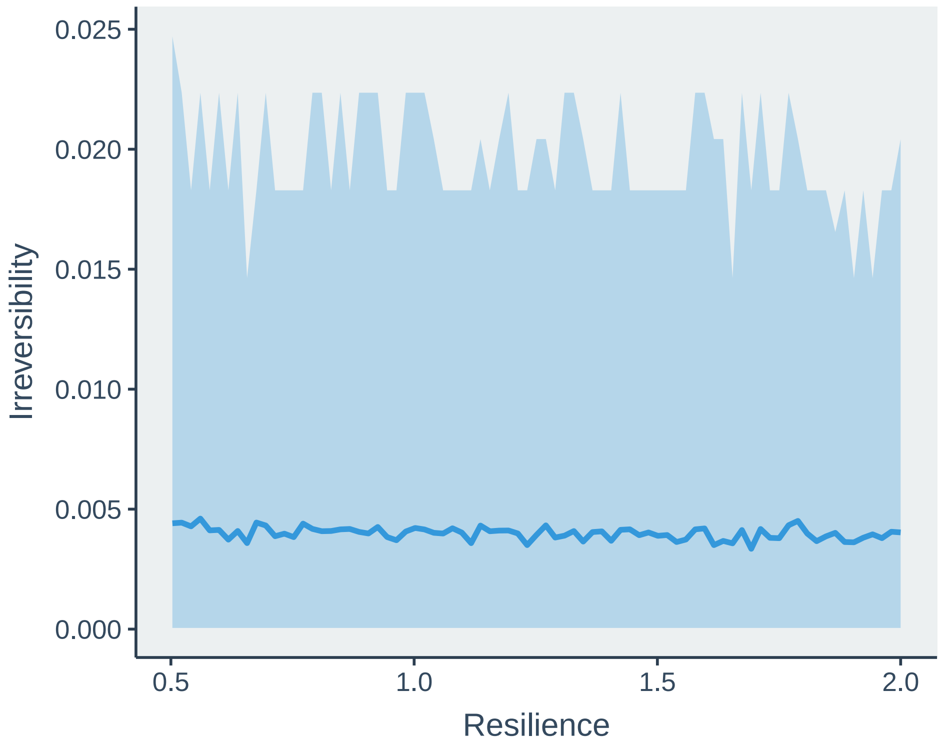


Figure S9: Association between irreversibility and resilience in logistic growth model. The line denotes the mean, and the ribbon denotes the 95% confidence interval.

#### E.2 Proof for multiple species with random interactions

To link with the theory of fluctuations in Lotka-Volterra dynamics, we consider the multispecies Lotka-Volterra dynamics,

$$\frac{\mathrm{d}\mathbf{N}}{\mathrm{d}t} = \mathrm{diag}(\mathbf{N})(\mathbf{r} + \mathbf{A}\mathbf{N}) \tag{S53}$$

Lotka-Volterra dynamics are scale-invariant. Without loss of generality, we can assume the equilibrium abundance is $\mathbf{N}^* = \mathbf{1}$ (a vector of ones) by scaling the growth rates $\mathbf{r}$ appropriately.

We now focus on random interaction matrices to obtain analytical results. We assume that the intraspecific interaction are all $\mu_d < 0$, and the interspecific interactions $A_{ij}$ are drawn independently from a distribution with mean $\mu > 0$ and variance $\sigma^2$.

Under this assumption, the resilience at equilibrium $\lambda^*$ (the largest real part of the eigenvalues of $\mathbf{J}$ at equilibrium) is:

$$\lambda^* = \mu_d + (S-1)\mu \tag{S54}$$

where $S$ is the number of species.

Now consider the system out of equilibrium. The time-varying Jacobian $\mathbf{J}$ is

$$\mathbf{J} = \frac{\partial \frac{\mathrm{d}\mathbf{N}}{\mathrm{d}t}}{\partial \mathbf{N}} = \mathrm{diag}(\mathbf{N})\mathbf{A} \tag{S55}$$

We consider the long-term average dynamical behaviors. A fundamental property of Lotka-Volterra systems is that the long-term average of the species abundances must equal the equilibrium abundances (see Theorem 5.2.3 in [8]). Thus, we can consider that $\mathbf{N}$ is generated from an arbitrary distribution with positive support with mean 1 and variance $\sigma_N^2$. The long run average Jacobian identical to the Jacobian at equilibrium, and hence (following the results in [9]):

$$\overline{\lambda} = \mu_d + (S-1)\mu = \lambda^* \tag{S56}$$

Thus, the long-term average resilience in a Lotka-Volterra system with random mutualistic interactions is identical to the resilience at equilibrium. This holds regardless of the magnitude of the fluctuations around the equilibrium. This result highlights the disconnect between resilience (a measure of stability) and irreversibility, as the latter can change with fluctuations even when the former remains constant.

### F Theoretical analysis in predation dynamics

#### F.1 Lotka-Volterra predation dynamics

The classic Lotka-Volterra (LV) predation model describes the coupled dynamics of prey ($N$) and predator ($P$) populations:

$$\frac{1}{N}\frac{\mathrm{d}N}{\mathrm{d}t} = \alpha - \beta P, \tag{S57}$$

$$\frac{1}{P}\frac{\mathrm{d}P}{\mathrm{d}t} = \delta N - \gamma. \tag{S58}$$

To simplify the analysis without altering the system's irreversibility, we rescale the variables and parameters:

$$\tilde{N} = \frac{\gamma}{\delta}N, \quad \tilde{P} = \frac{\gamma}{\beta}P, \text{and} \quad \tilde{t} = \frac{1}{\gamma}t, \tag{S59}$$

leading to the rescaled LV equations:

$$\frac{1}{\tilde{N}}\frac{\mathrm{d}\tilde{N}}{\mathrm{d}\tilde{t}} = A - \tilde{P}, \tag{S60}$$

$$\frac{1}{\tilde{P}}\frac{\mathrm{d}\tilde{P}}{\mathrm{d}\tilde{t}} = \tilde{N} - 1. \tag{S61}$$

where $A = \alpha/\gamma$.

The LV model predicts perpetual cycles in prey and predator abundances, as illustrated in Figure S10. The cycles become more asymmetric with larger perturbation:

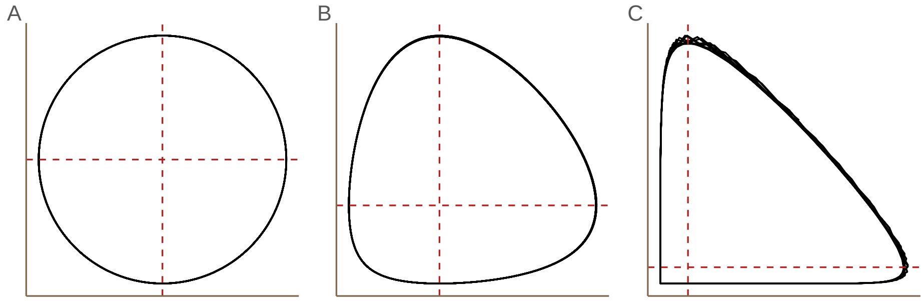


Figure S10: Low, medium and high irreversibility

The Lotka-Volterra predation dynamics is Hamiltonian with Hamiltonian function

$$H := N - \log N + P - A\log P \tag{S62}$$

To determine the period, we first find the maximum and minimum values of N (corresponding to $\frac{\mathrm{d}N}{\mathrm{d}t} = 0$, and $P^* = A$). Substituting this into the Hamiltonian yields

$$N - \log N = A - A\log A - H \tag{S63}$$

Solving this equation gives the minimum and maximum values of $N$:

$$N_{\min} = -W_0\left[-\frac{e^{A-H}}{A^A}\right], \tag{S64}$$

$$N_{\max} = -W_{-1}\left[-\frac{e^{A-H}}{A^A}\right], \tag{S65}$$

where $W_k$ denotes the $k$-th branch of the Lambert W function [10].

We then express $P$ as a function of $N$. With similar technique, we obtain $P$ for the upper and lower branches:

$$P_{\mathrm{upper}} = -AW_{-1}\left[-\frac{1}{A}N^{-1/A}e^{(N-H)/A}\right], \tag{S66}$$

$$P_{\mathrm{lower}} = -AW_0\left[-\frac{1}{\alpha}N^{-1/A}e^{(N-H)/A}\right]. \tag{S67}$$

The irreversibility $I$ then is given by

$$I = \frac{1}{t_{\uparrow\uparrow}+t_{\uparrow\downarrow}+t_{\downarrow\uparrow}+t_{\downarrow\downarrow}}\left((t_{\uparrow\uparrow}-t_{\downarrow\downarrow})\log\frac{t_{\uparrow\uparrow}}{t_{\downarrow\downarrow}}+(t_{\uparrow\downarrow}-t_{\downarrow\uparrow})\log\frac{t_{\uparrow\downarrow}}{t_{\downarrow\uparrow}}\right) \tag{S68}$$

where the period lengths for each phase of the cycle ($\Omega = \{\uparrow\uparrow, \uparrow\downarrow, \downarrow\uparrow, \downarrow\downarrow\}$) are [11]

$$t_{\uparrow\uparrow} = \int_1^{N_{\max}} \frac{1}{AN(1+P_{\mathrm{lower}})}dN, \tag{S69}$$

$$t_{\downarrow\uparrow} = \int_{N_{\max}}^1 \frac{1}{AN(1+P_{\mathrm{upper}})}dN, \tag{S70}$$

$$t_{\downarrow\downarrow} = \int_{N_{\min}}^1 \frac{1}{AN(1+P_{\mathrm{upper}})}dN, \tag{S71}$$

$$t_{\uparrow\downarrow} = \int_1^{N_{\min}} \frac{1}{AN(1+P_{\mathrm{lower}})}dN. \tag{S72}$$

While a closed-form expression for $I$ remains elusive, simulations (Figure S11) clearly show that increasing $\alpha$ (i.e., increasing the ecosystem's energy) amplifies irreversibility.

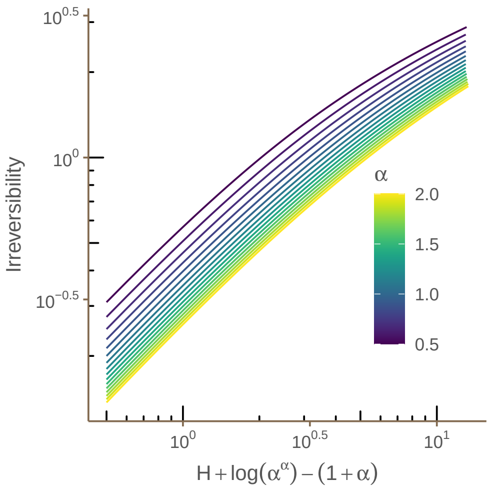


Figure S11: Ecosystem energy dominates irreversibility.

Furthermore, we developed a prediction (Figure S12) based on these simulations.

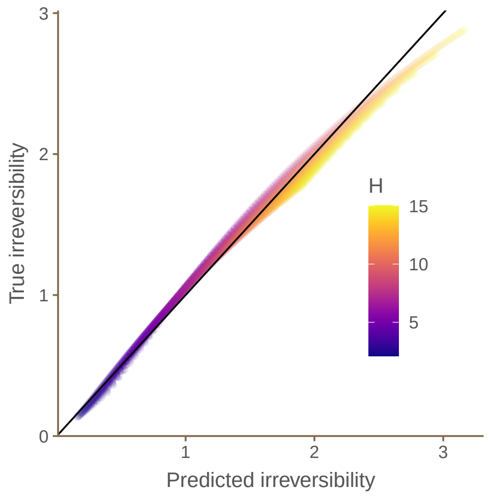


Figure S12: Prediction

#### F.2 Conservative ecological dynamics

We now generalize to more than 2 species. In this section, we focus on conservative ecological dynamics. Specifically, a conservative Lotka-Volterra dynamics is defined as a system with interaction matrix $A$ where there exists a matrix $D > 0$ such that $AD$ is skew-symmetric. It has been proved that a conservative Lotka-Volterra dynamics always admits a Hamiltonian [12].

##### F.2.1 3 species food chain

We consider the dynamics of a 3 species food chain

$$\frac{\mathrm{d}x_1}{\mathrm{d}t} = x_1 \left( r_1 + \omega_1 x_2 - \omega_2 x_3 \right) \tag{S73}$$

$$\frac{\mathrm{d}x_2}{\mathrm{d}t} = x_2 \left(r_2 - \omega_1 x_1 + \omega_3 x_3\right) \tag{S74}$$

$$\frac{\mathrm{d}x_3}{\mathrm{d}t} = x_3 \left(r_3 + \omega_2 x_1 - \omega_3 x_2\right) \tag{S75}$$

The Hamiltonian of this system is given by

$$H = x_1 + x_2 + x_3 - \frac{r_2}{\omega_1} \log x_1 + \frac{r_1}{\omega_1} \log x_2. \tag{S76}$$

With simulations, we found again a strong positive association between energy and irreversibility.

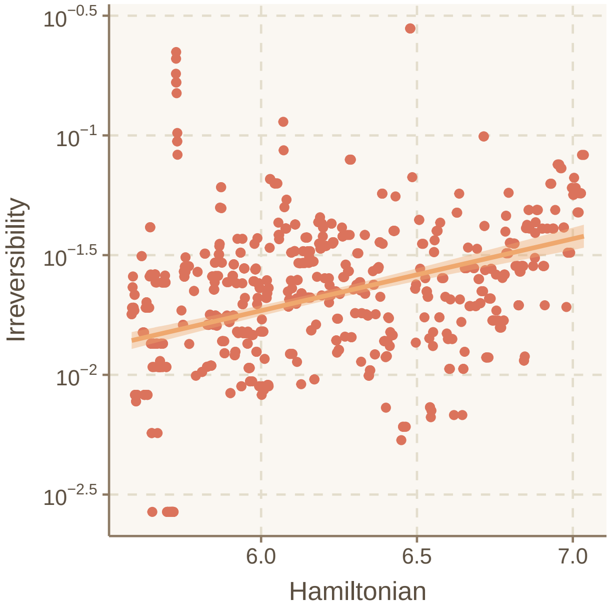


Figure S13: Positive association between Hamiltonian and irreversibility in 3 species food chain.

##### F.2.2 4 species food web

Similarly, we can consider the dynamics of a 4-species food web:

$$\frac{\mathrm{d}x_1}{\mathrm{d}t} = x_1 \left(-1 + x_2\right) \tag{S77}$$

$$\frac{\mathrm{d}x_2}{\mathrm{d}t} = x_2 \left(1 - x_1 + a x_3\right) \tag{S78}$$

$$\frac{\mathrm{d}x_3}{\mathrm{d}t} = x_3 \left(-1 - a x_2 + x_4\right) \tag{S79}$$

$$\frac{\mathrm{d}x_4}{\mathrm{d}t} = x_4 \left(1 - x_3\right) \tag{S80}$$

The Hamiltonian of this system is given by

$$H = x_1 + x_2 + x_3 + x_4 - (1 + a) \log \left(x_1 x_4\right) - \log \left(x_2 x_3\right) \tag{S81}$$

With simulations, we found again a strong positive association between energy and irreversibility.

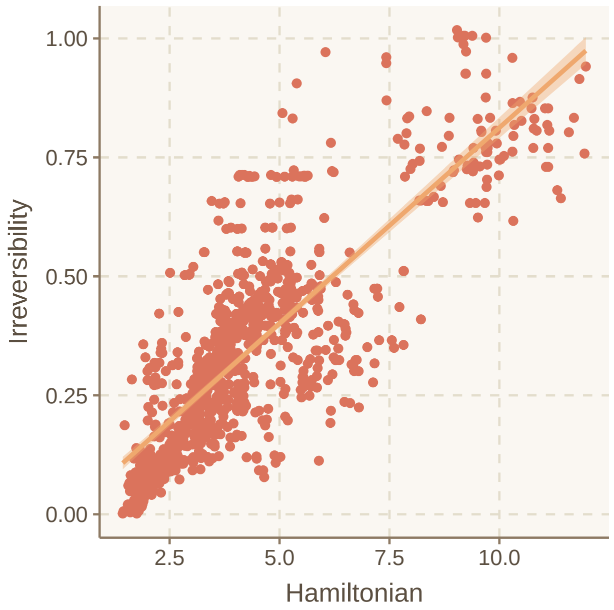


Figure S14: Positive association between Hamiltonian and irreversibility in 4 species food web.

##### F.3 Linearized Lotka-Volterra predation dynamics

It is important to note that the linearized LV model is completely reversible. The linearized LV dynamics reads as [13, 14]

$$\frac{\mathrm{d}N}{\mathrm{d}t} = -\frac{bd}{c}\left(P - \frac{a}{b}\right) \tag{S82}$$

$$\frac{\mathrm{d}P}{\mathrm{d}t} = \frac{ac}{b}\left(N - \frac{d}{c}\right) \tag{S83}$$

Their trajectory has an analytical solution, forming an ellipse (see Figure S15):

$$N(t) = \frac{d}{c} + r\sin\left(t\sqrt{ad} + t_*\right) \tag{S84}$$

$$P(t) = \frac{a}{b} + r\frac{c}{b}\sqrt{\frac{a}{d}}\cos\left(t\sqrt{ad} + t_*\right) \tag{S85}$$

where $t^*$ satisfies the following conditions:

$$\cos(t_*) = \frac{P_0 - a/b}{r}\frac{b}{c}\sqrt{\frac{d}{a}} \tag{S86}$$

$$\sin(t_*) = \frac{N_0 - d/c}{r}, \tag{S87}$$

$$r = \sqrt{\left(u_0 - \frac{d}{c}\right)^2 + \left(v_0 - \frac{a}{b}\right)^2 \frac{b^2 d}{ac^2}} \tag{S88}$$

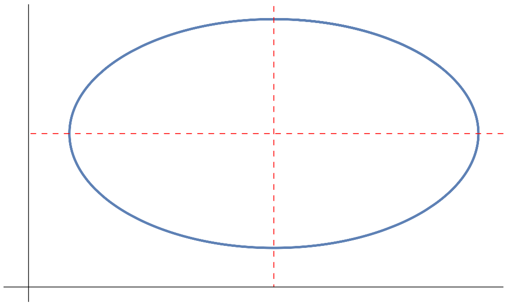

Figure S15

In this linearized model, the forward and reverse trajectories have identical probabilities, due tot the symmetry in the ellipse trajectory. For example, $\mathcal{F}_{\uparrow\uparrow} = \mathcal{F}_{\downarrow\downarrow} = \mathcal{R}_{\uparrow\uparrow}$. Thus, it is always completely reversible.

### G Empirical analysis in predation dynamics

#### G.1 Empirical analysis on temporal fluctuation

To empirically investigate the relationship between irreversibility and fluctuations in predator-prey dynamics, we utilized a high-quality, long-term time series dataset [15]. We focused on the five replicates (experimental labels C1-C5) as they have identical experimental setups. The predator is *Brachionus calyciflorus* and the prey is *Monoraphidium minutum*. The environmental conditions are constant: dilution rate $\delta = 0.55$/day, and concentration of the external medium Nin $= 80$ $\mu$mol/L. Figure S16 shows the time series.

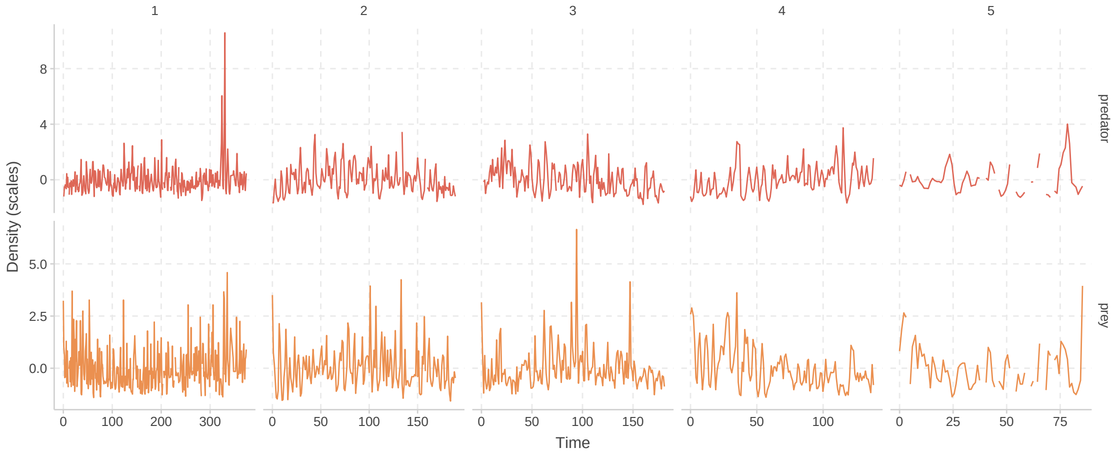


Figure S16: Time series of prey (green) and predator (orange) abundances in the five replicates (C1-C5) of the planktonic predator-prey system experiment from [15].

Additionally, Figure S17 illustrates the phase plot of prey and predator abundances, showcasing the cyclical nature of their interactions.

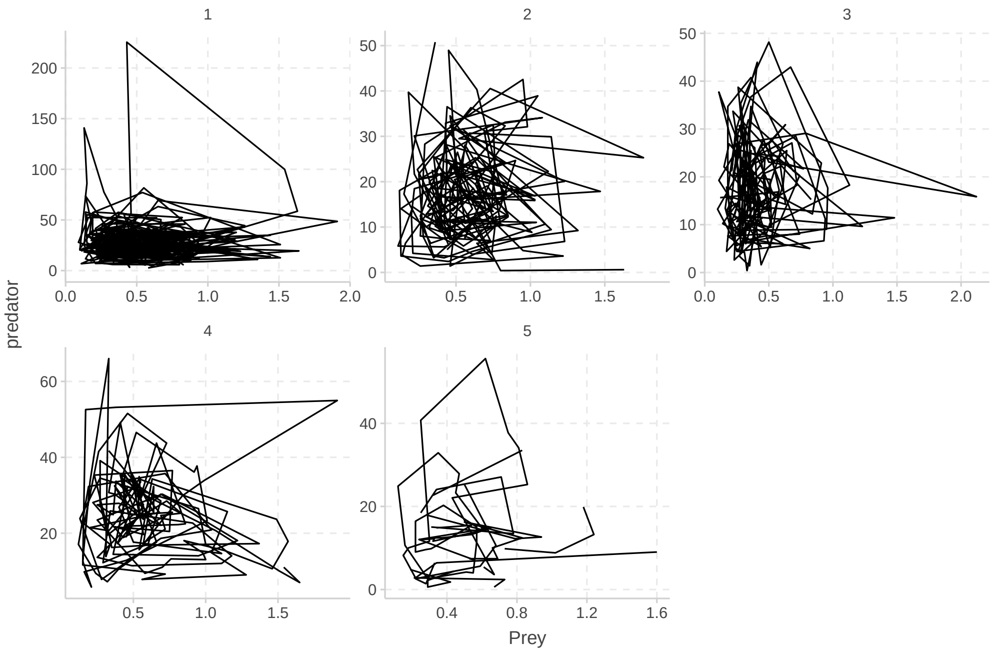


Figure S17: Phase plot of prey and predator abundances in the planktonic predator-prey system experiment from [15].

We quantify the minimum distance to equilibrium as the minimum Euclidean distance of empirical abundances from the mean abundance. The theoretical justification is that, in both Lotka-Volterra dynamics and the linearized dynamics, the equilibrium abundance is the same as the average abundance throughout the trajectory. Note that this distance is generally not comparable across different systems. However, here we are studying the same empirical system with identical conditions, allowing for comparison.

Figure 2D in the main text presents the average relationship between distance from equilibrium and irreversibility. To provide a more comprehensive analysis, we conducted a bootstrapping analysis to assess the robustness of this relationship. The results of this analysis are shown in Figures S18 and S19, with Figure S18 displaying the aggregated results and Figure S19 presenting the results for each individual replicate. The positive association between distance from equilibrium and irreversibility is evident in both figures, supporting the hypothesis that fluctuations away from equilibrium lead to increased irreversibility in predator-prey dynamics.

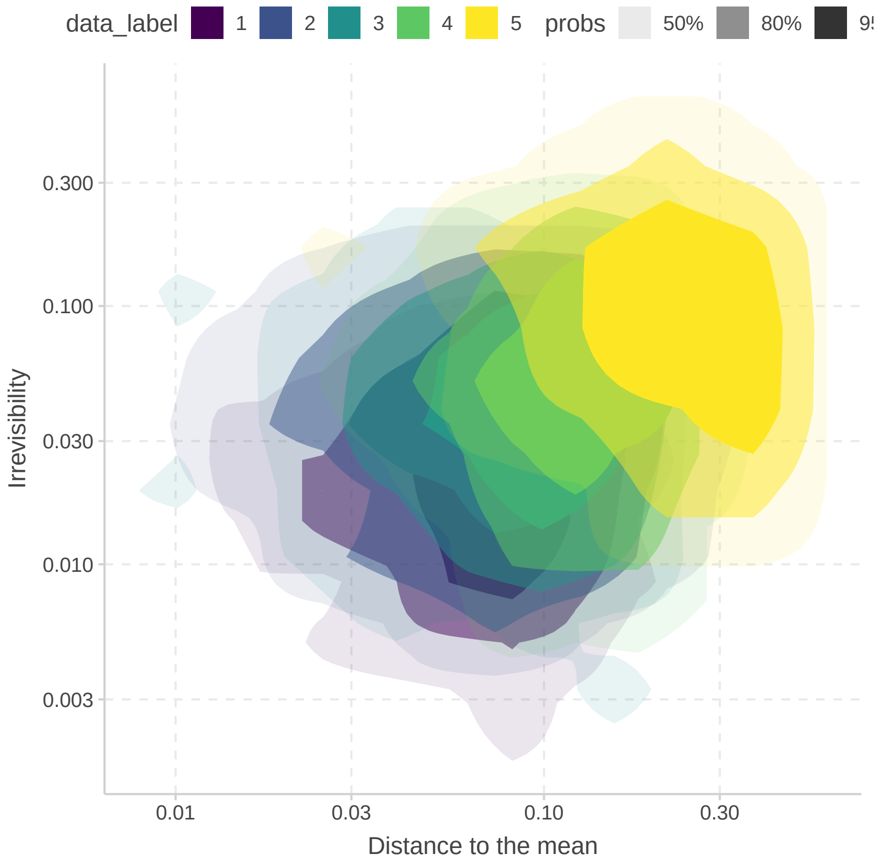


Figure S18: Bootstrapped estimates of the relationship between minimum distance to equilibrium and irreversibility in the planktonic predator-prey system. The shaded area shows the confidence level, and the color corresponds to different replicates.

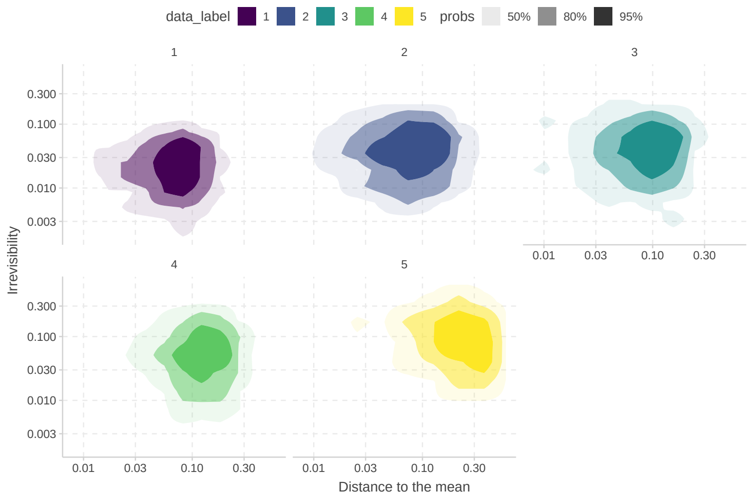


Figure S19

#### G.2 Empirical analysis on rapid evolution

To investigate the impact of rapid evolution on consumer-resource dynamics, we analyzed two distinct groups of time series data. For consumer-resource dynamics without rapid evolution, we analyzed 18 time series across diverse ecosystems [15, 16, 17, 18, 19, 20, 21, 22], compiled and processed by [23] except for [15]. Figure S20 shows their time series.

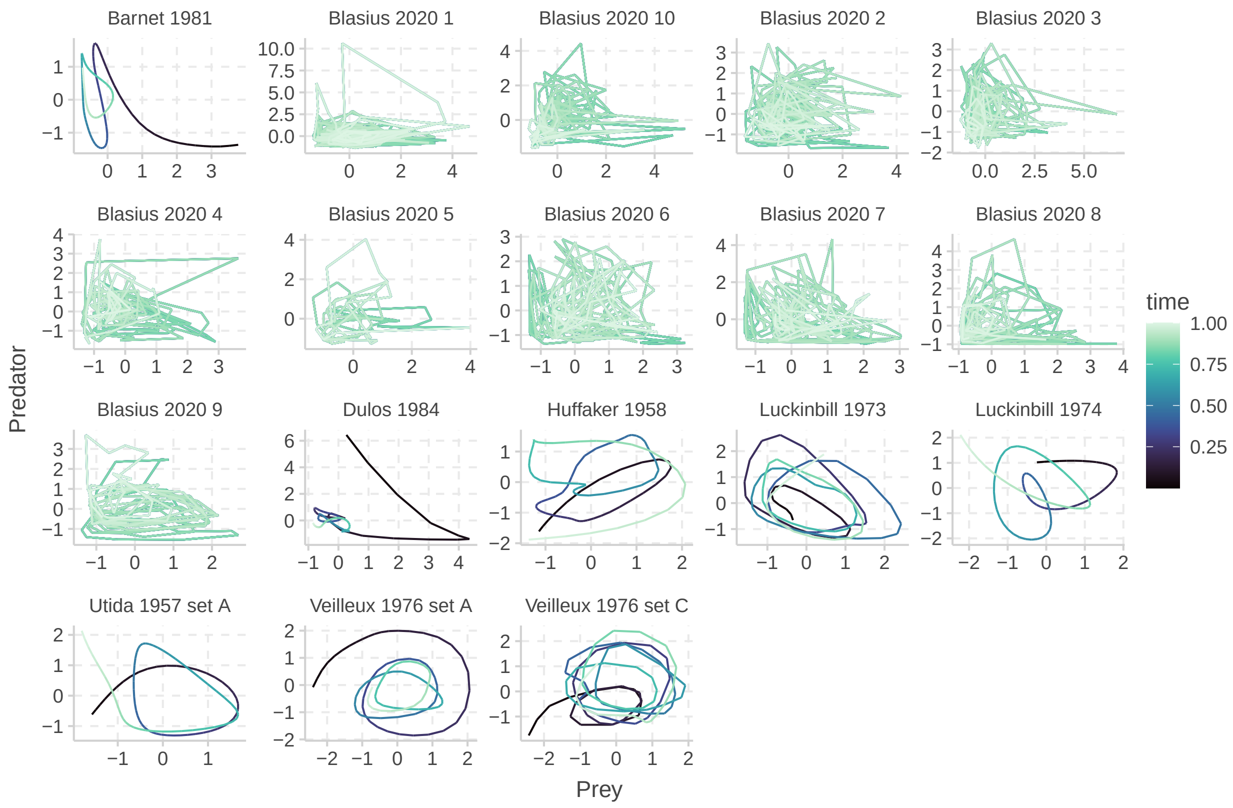


Figure S20: Time series of consumer-resource dynamics across diverse ecosystems with little evidence of rapid evolution.

To compare, we analyzed 13 prey-predator time series where rapid evolution is observed [16, 22, 24, 25, 26, 27, 28, 29, 30, 31]. These datasets, compiled and processed by [23], encompass a diverse range of ecosystems, providing an ideal testbed. Figure S21 shows their time series.

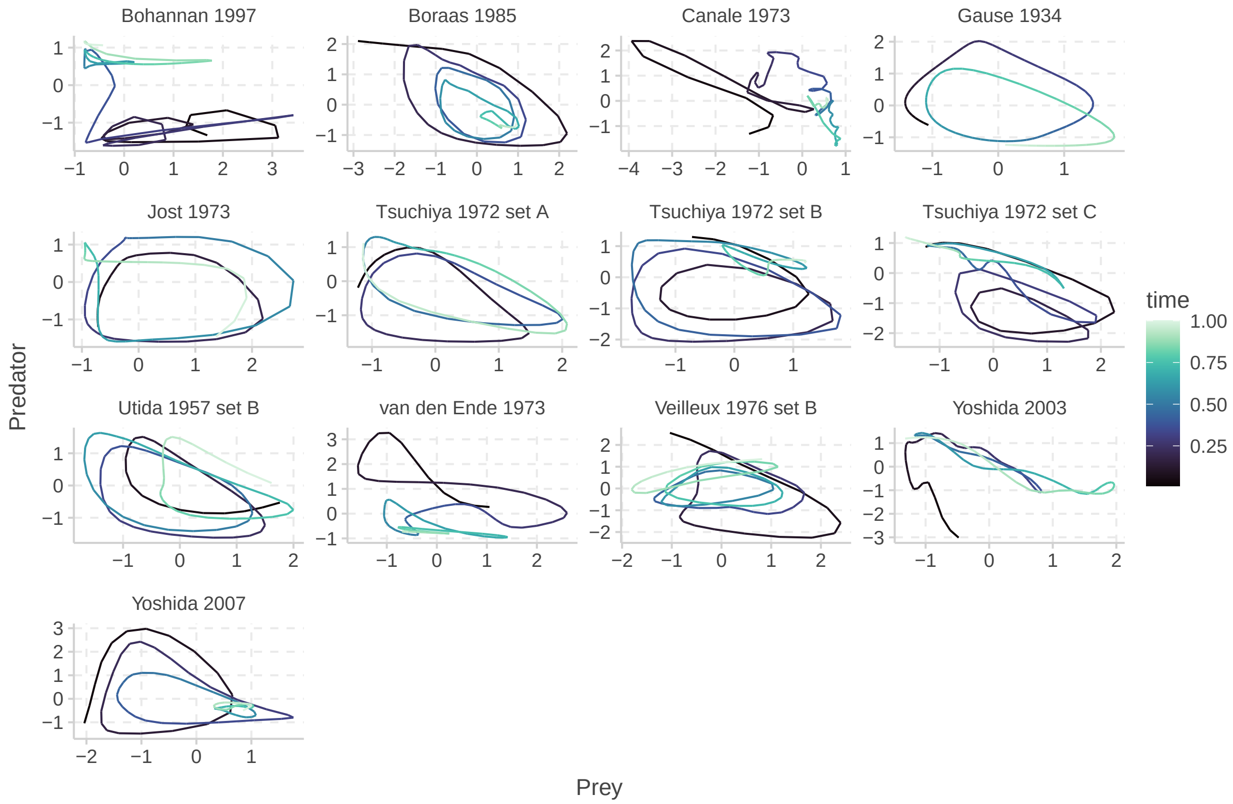


Figure S21: Time series of consumer-resource dynamics across diverse ecosystems with strong evidence of rapid evolution.

A key distinction between these two groups lies in the stationarity of their dynamics. Systems without rapid evolution tend to exhibit stationary dynamics, while those with rapid evolution often display non-stationary dynamics. This non-stationarity arises because evolution introduces new equilibria, shifting the system's dynamics over time.

To rigorously assess the stationarity of the time series, we employed the Augmented Dickey-Fuller (ADF) test. This test examines the presence of a unit root, which is indicative of non-stationarity. Time series with a trend, as often observed in systems with rapid evolution, typically possess a unit root and yield a large p-value in the ADF test (Figure S22).

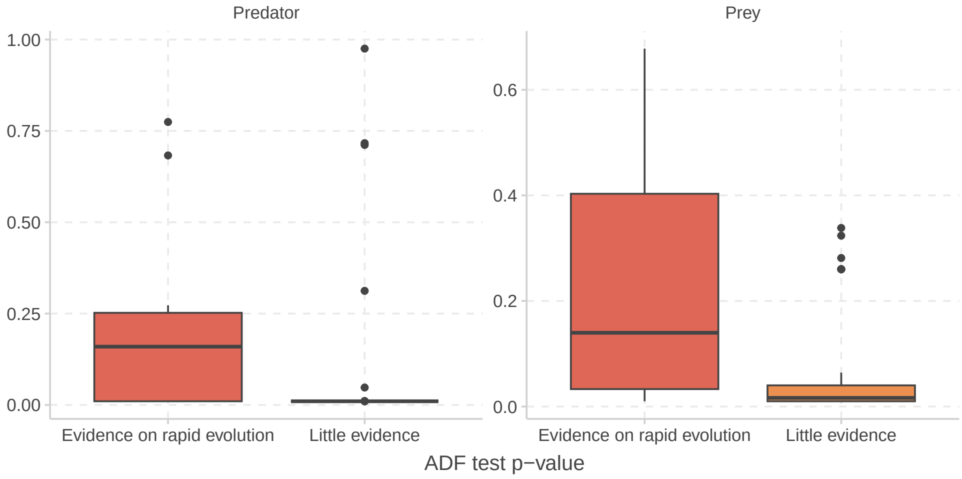


Figure S22: ADF test results for consumer-resource dynamics with and without rapid evolution. The p-values indicate the significance of the test for a unit root, with larger p-values suggesting non-stationarity.

Bearing this in mind, we now study the irreversibility of these systems. We conducted a convergence analysis to examine how irreversibility changes over time (Figure S23).

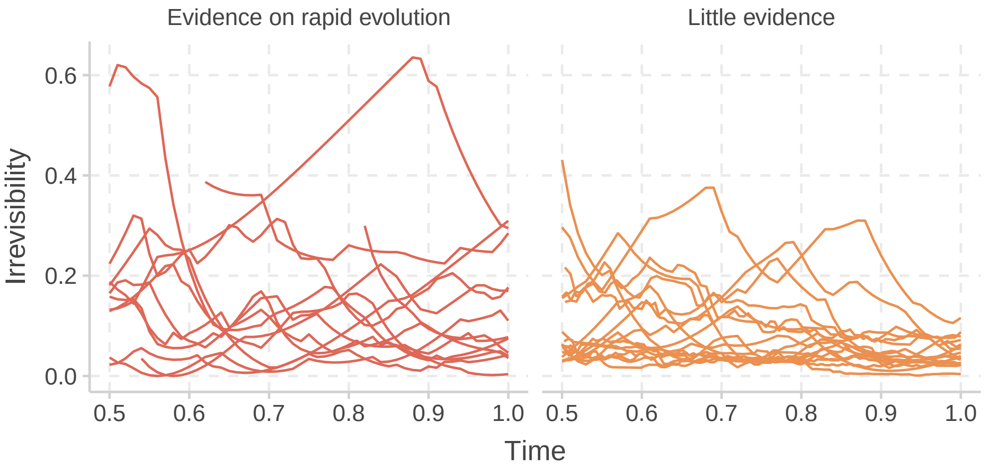


Figure S23: Convergence analysis of irreversibility for consumer-resource dynamics with and without rapid evolution. The plots show how irreversibility changes over time for each time series.

Visual inspection of Figure S23 reveals a clear pattern: irreversibility tends to converge in systems without rapid evolution, while it often diverges or remains non-convergent in systems with rapid evolution. This observation is further supported by Figure S24.

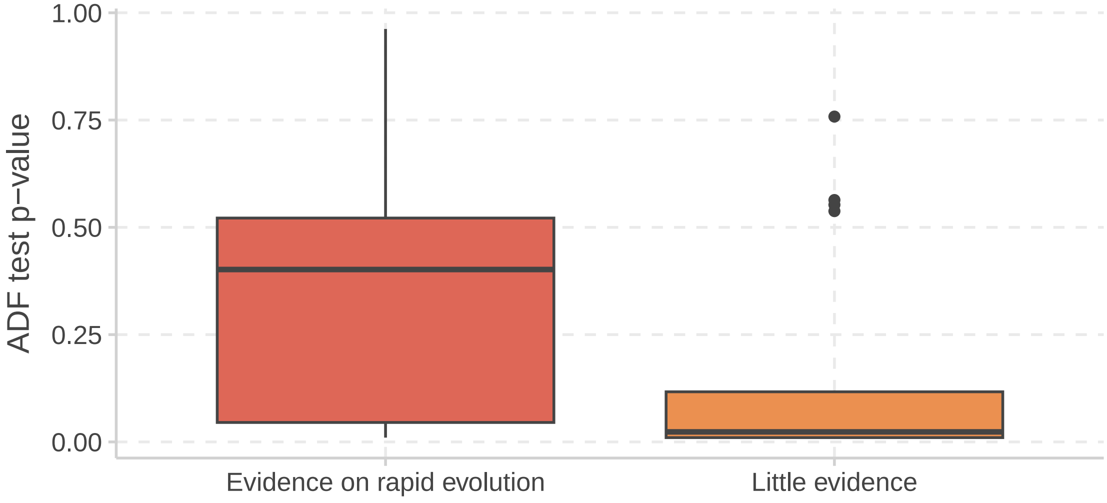


Figure S24: Stationarity of irreversibility in consumer-resource dynamics with and without rapid revolution.

In conclusion, our analysis of empirical data suggests that rapid evolution plays a crucial role in shaping the irreversibility of consumer-resource dynamics. Systems with rapid evolution tend to exhibit non-stationary dynamics and higher irreversibility, while those without rapid evolution tend to exhibit stationary dynamics and lower irreversibility.

### H Theoretical analysis in multispecies dynamics with unstructured interaction

We consider stochastic Lotka-Volterra dynamics with dispersal,

$$\mathrm{d}N_i = N_i(r_i - \sum_j a_{ij}N_j)\mathrm{d}t + D\mathrm{d}t + \eta N_i \mathrm{d}W \tag{S89}$$

where $N_i$ represents the abundance of species i, $a_{ij}$ is the competition coefficient between species $i$ and $j$ (drawn from some distribution), $D$ is the dispersal rate, $\eta$ is the noise level, and $\mathrm{d}W$ denotes differential form of the Brownian motion.

It is well-established that the dynamics of systems with random species interactions exhibit three distinct phases [32, 33]:

- **Stable Phase**: Characterized by low fluctuations in species abundances, primarily driven by stochastic noise (e.g., demographic stochasticity or environmental fluctuations). In this phase, species interactions are relatively weak, and the system tends to remain close to its equilibrium state.

- **Oscillatory Phase**: Characterized by regular or quasi-regular oscillations in species abundances. These oscillations arise from the interplay between species interactions and stochasticity. The amplitude and frequency of the oscillations depend on the strength of species interactions and the level of noise.

- **Irregular/Chaotic Phase**: Characterized by large, irregular fluctuations in species abundances, potentially exhibiting chaotic behavior. In this phase, species interactions are strong, leading to complex and unpredictable dynamics. The system may exhibit sensitivity to initial conditions and long-term unpredictability.

These phases are universal, meaning they occur across a wide range of parameter values and functional forms of species interactions. The transitions between phases are primarily determined by two key factors: species richness (the number of species in the community) and the average strength of species interactions. Increasing either of these factors tends to drive the system from the stable phase to the oscillatory phase and eventually to the irregular/chaotic phase.

Following [34], we sample non-diagonal $a_{ij}$ from a uniform distribution $[0, 2\alpha]$, and parametrize the diagonal as $a_{ii} = 1$ and the $r_i$ as 1. Figure 3A shows the general phase plot with average behavior, and clearly exhibits these three phases. Figure S25 he variation in dynamics across different parameter combinations.

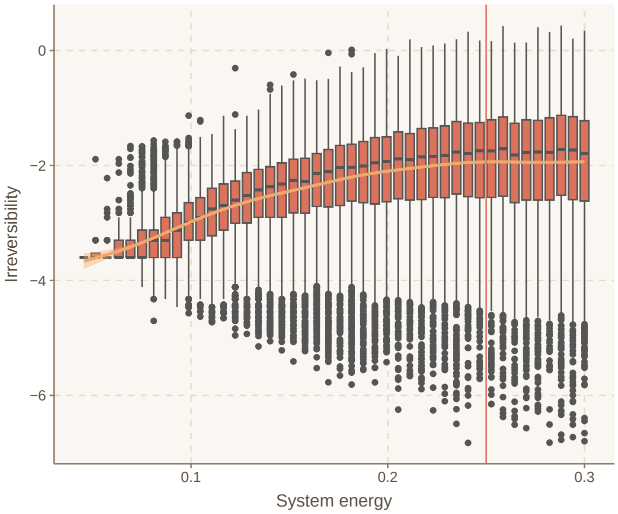


Figure S25

Our analysis suggests that the phase transition thresholds in this model are more aligned with May's classic result [35], which posits an identical role for species richness and average competition strength in determining the dynamical regime. This contrasts slightly with the phase transition criteria in [32], although the general trend remains similar. This discrepancy warrants further investigation in future studies.

### I Empirical analysis in multispecies dynamics with unstructured interaction

We tested our predictions using microbiome data [34]. The authors manipulated species pool size and nutrient availability (which affects species interactions) to examine fluctuation dynamics. They measured both biomass and abundance, but we exclusively used the abundance data as biomass data was only available as community-level aggregates, unsuitable for our analysis. Their analysis suggested that both measures yield similar results.

Figure S26 illustrates the time series for different species richness and nutrient levels.

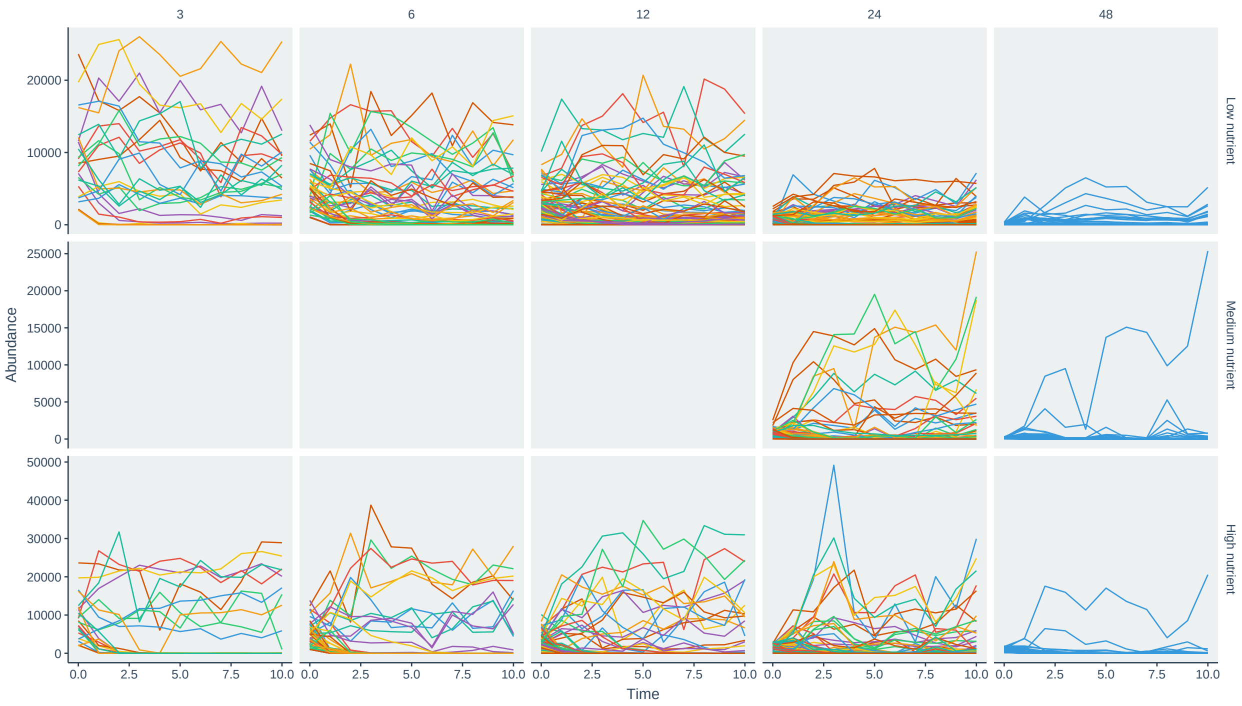


Figure S26: Time series of species abundances in microbiome communities with varying species richness (columns) and nutrient levels (rows). Different colors correspond to different species.

Figure 2C in the main text presents the results for low nutrient conditions. S27 compares low and high nutrient conditions across a gradient of species richness. As predicted by theory (Figure 2A), high nutrient conditions consistently exhibit the most stable irreversibility, as it is in the high fluctuations phase even with low species richness.

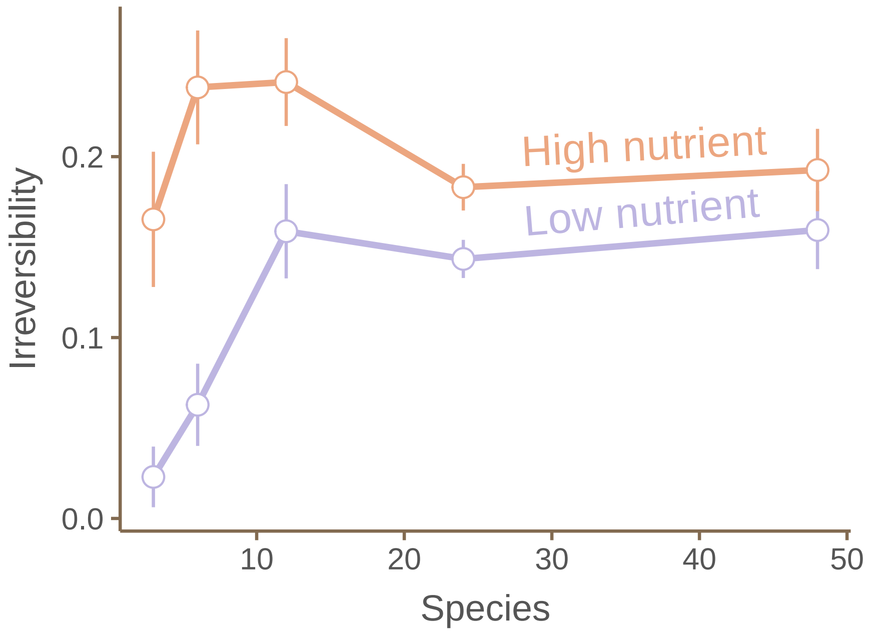


Figure S27: Comparison of irreversibility between low (purple) and high (orange) nutrient conditions across a gradient of species richness in microbiome communities. Points represent the mean irreversibility, and error bars represent the 95% confidence interval.

To examine the saturation effect of high species richness, Figure S28 further demonstrates the statistical indistinguishability of irreversibility at high species richness levels.

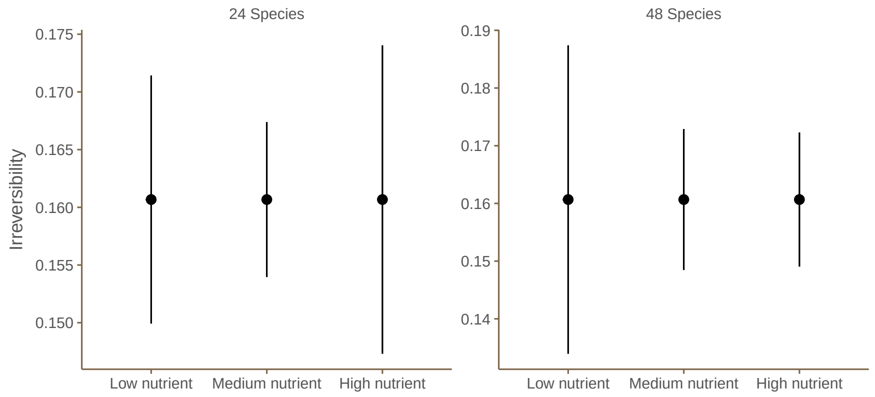


Figure S28

Due to the limited number of data points (10) in each time series, which is insufficient for accurate irreversibility estimation, we employed a standard time series analysis technique using a Gaussian kernel to smooth the data. We tested the robustness of the observed patterns by considering various combinations of hyperparameters for bandwidth and smoothed points. The mean standard deviation of the irreversibility estimates across these combinations was consistently below 2.5%, indicating the robustness of our results to smoothing parameters (Figure S29).

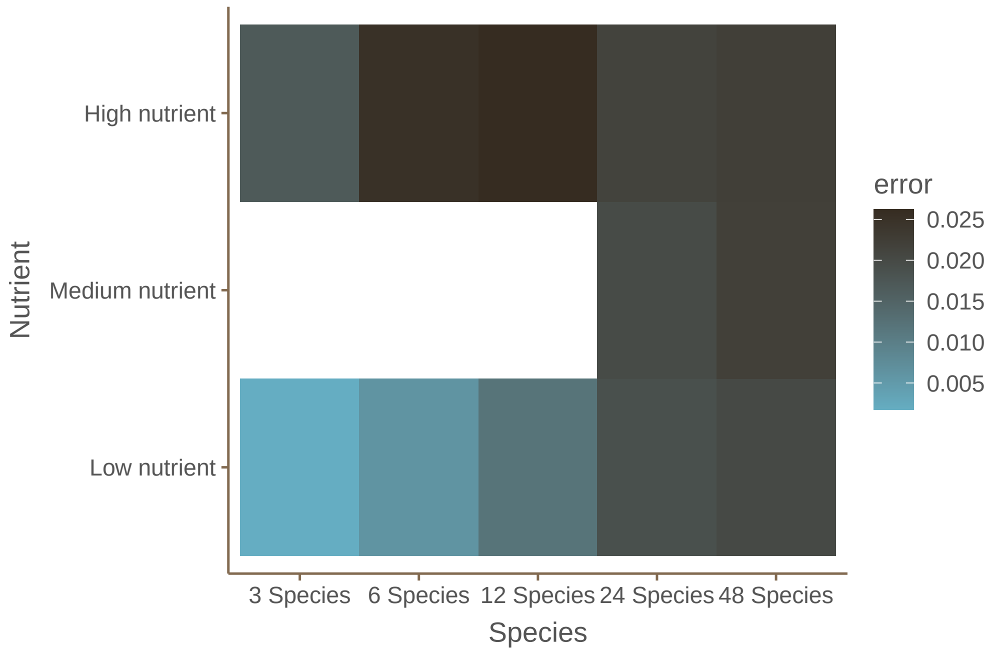
High nutrient
Medium nutrient
Low nutrient
Nutrient
3 Species
6 Species
12 Species
24 Species
48 Species
Species
error
0.025
0.020
0.015
0.010
0.005

Figure S29

### J Theoretical analysis in multispecies dynamics with metabolic constraints

In the main text, we consider a Lotka-Volterra competition model constrained by metabolic relationships to investigate the interplay between metabolic constraints and ecological dynamics. Here, we consider more general relationships between metabolic rates, body mass, and temperature.

Specifically, we assume that the intrinsic growth rates ($r_i$) and competition coefficients ($a_{ij}$) scale with body mass ($M$) and temperature ($T$) according to the following equations [36]:

$$r_i \propto e^{-E_i/k_\mathrm{B}T} \cdot M_i^\beta \tag{S90}$$

$$a_{ij} \propto \cdot e^{\left(E_i - E_j\right)/k_\mathrm{B}T} \cdot \left(M_i/M_j\right)^\alpha \tag{S91}$$

where $E_i$ is the activation energy of metabolism for species $i$, $k$ is the Boltzmann constant, and $\alpha$ and $\beta$ are scaling exponents. Metabolic scaling theory suggests that $\alpha = 3/4$ and $\beta = -3/4$ across many species, although alternative values like $\alpha = -1$ and $\beta = 1$ have been observed in certain bioregimes [37]. Notably, these scaling relationships maintain the constraint $\alpha + \beta = 0$.

Figure 4B in the main text has shown that irreversibility decreases with higher temperature with exponent $\alpha = 3/4$. Here, we further show that it holds for other $\alpha$s (Figure S30):

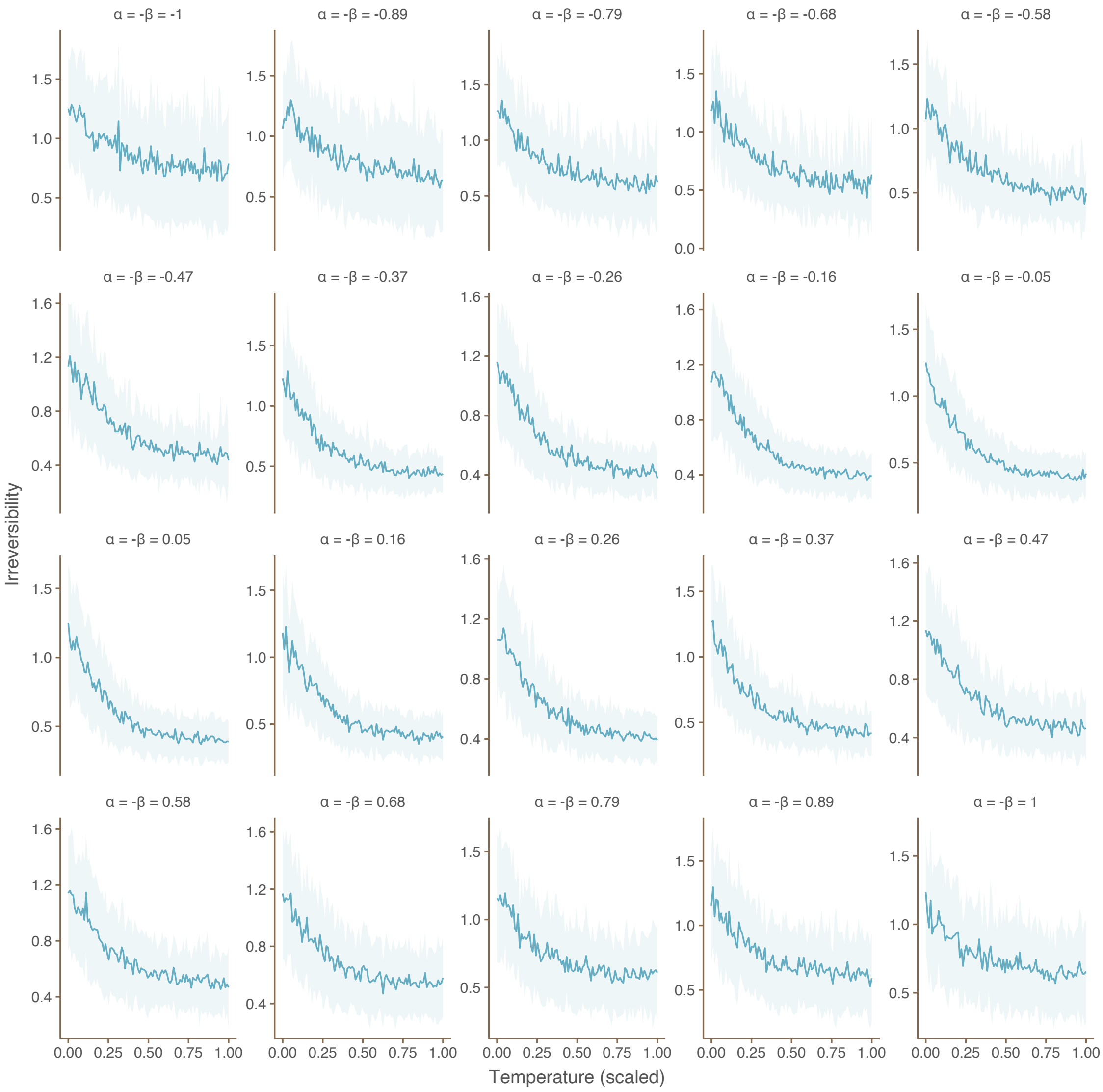


Figure S30: Irreversibility decreases with temperature under different metabolic scaling exponents ($\alpha$). Each panel corresponds to a different value of $\alpha$, and the lines show the relationship between temperature and irreversibility for 100 replicate simulations. The negative slopes of the lines indicate that irreversibility consistently decreases with increasing temperature, regardless of the value of $\alpha$.

Furthermore, we show that irreversibility can predict the empirical pattern of $\alpha + \beta = 0$. To do so, we systematically varied the scaling exponents $\alpha$ and $\beta$ within a range of -2 to 2, computing all possible combinations. For each parameter combination, we simulated the Lotka-Volterra model and calculated the resulting irreversibility using otherwise identical setups.

Figure S31 illustrates the relationship between the scaling exponents ($\alpha$ and $\beta$) and irreversibility in the metabolically constrained Lotka-Volterra model. The results demonstrate that the lowest irreversibility values are consistently observed near the line where $\alpha + \beta = 0$, aligning with empirical observations across diverse taxa. This finding suggests that the empirical pattern of $\alpha + \beta = 0$ observed in nature may be a consequence of natural selection favoring systems with minimal irreversibility. This intriguing hypothesis warrants further investigation to elucidate the underlying mechanisms and evolutionary drivers behind this observed pattern.

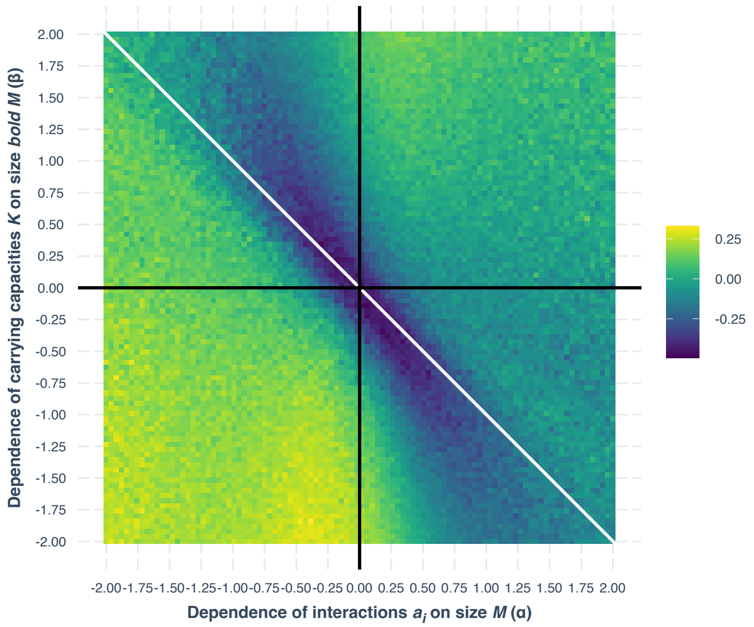


Figure S31: Relationship between scaling exponents (alpha and beta) and irreversibility in the metabolically constrained Lotka-Volterra model. The heatmap shows the irreversibility values for different combinations of alpha and beta, with warmer colors indicating higher irreversibility. The lowest irreversibility is achieved close to the line $\alpha + \beta = 0$.

### K Empirical analysis in multispecies dynamics with metabolic constraints

#### K.1 Empirical data analysis of an intertidal community

We analyzed a dataset of a rocky intertidal community in the Cape Rodney-Okakari Point Marine Reserve on the North Island of New Zealand [38]. It consisted of three sessile species: the honeycomb barnacle *Chamaesipho columna*, the crustosebrown alga *Ralfsia cf confusa*, and the little black mussel *Xenostrobus pulex*. Species abundances (expressedas percentage of cover) were monitored on a monthly basis for more than 20 years using a permanent grid. Figure S32 shows the time series of species abundance as well as the temperature.

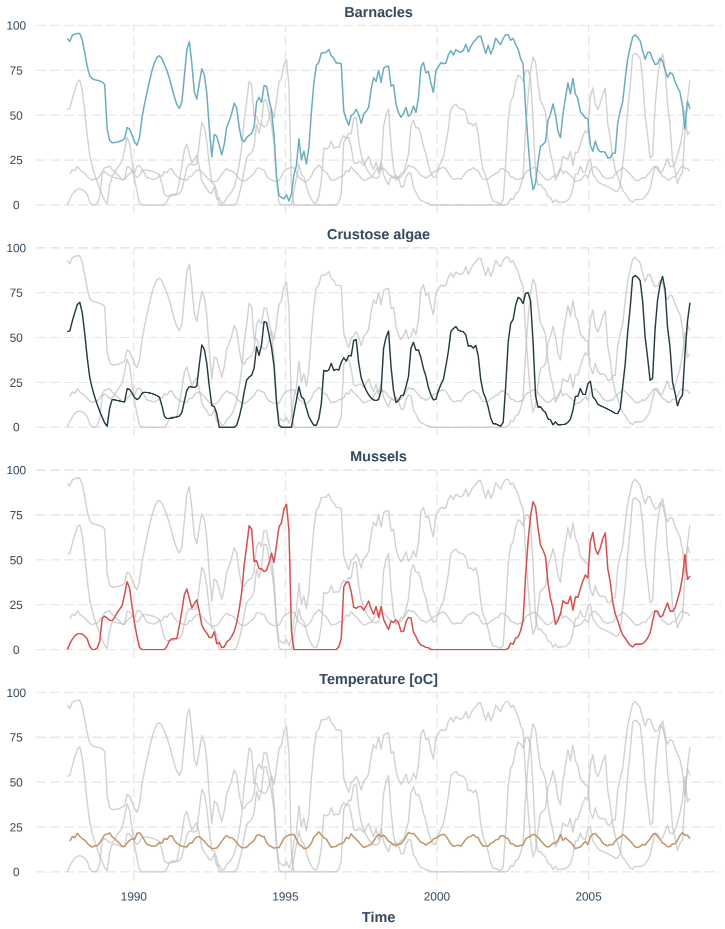


Figure S32: Time series of the intertidal community and the local temperature.

We first confirm whether irreversibility converges by gradually increasing observed time steps (Figure S33).

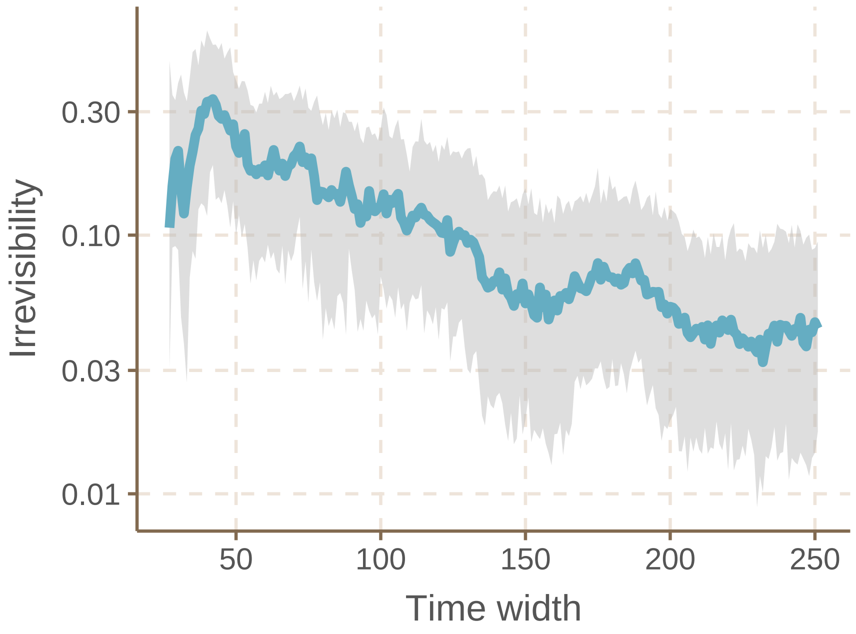


Figure S33: Convergence analysis

Figure 4C in the main text shows that temperature and irreversibility are negatively related. Figure S34 further validates the robustness of the conclusion by varying the length of the sliding window.

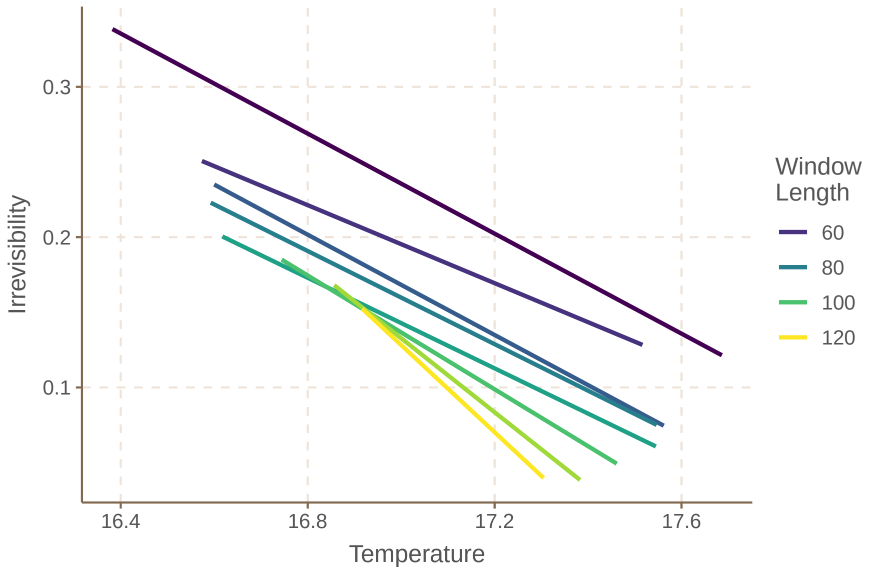


Figure S34: Generality of the negative association between mean temperature and irreversibility using varying window lengths.

#### K.2 Empirical data analysis of aquatic communities

We used five datasets of zooplankton and phytoplankton across various aquatic environments. Figure S35 shows the time series of species abundance as well as the temperature in the five time series (Lake Greifensee [39], Narragansett Bay [40], Wadden Sea [41, 42, 43]).

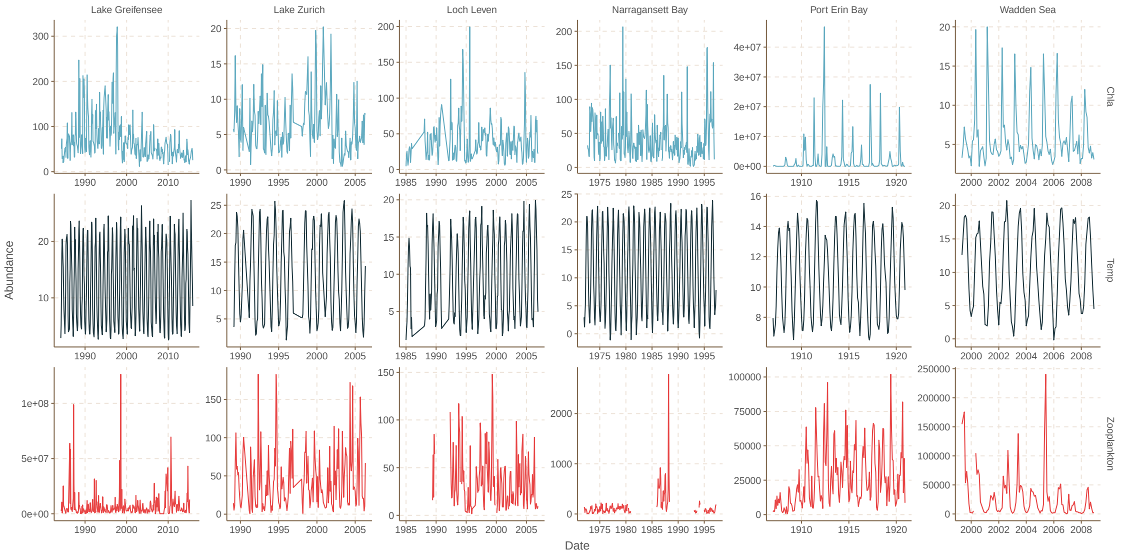


Figure S35: Time series of zooplankton and phytoplankton, and the local temperature.

We first confirm whether irreversibility converges by gradually increasing observed time steps (Figure S36).

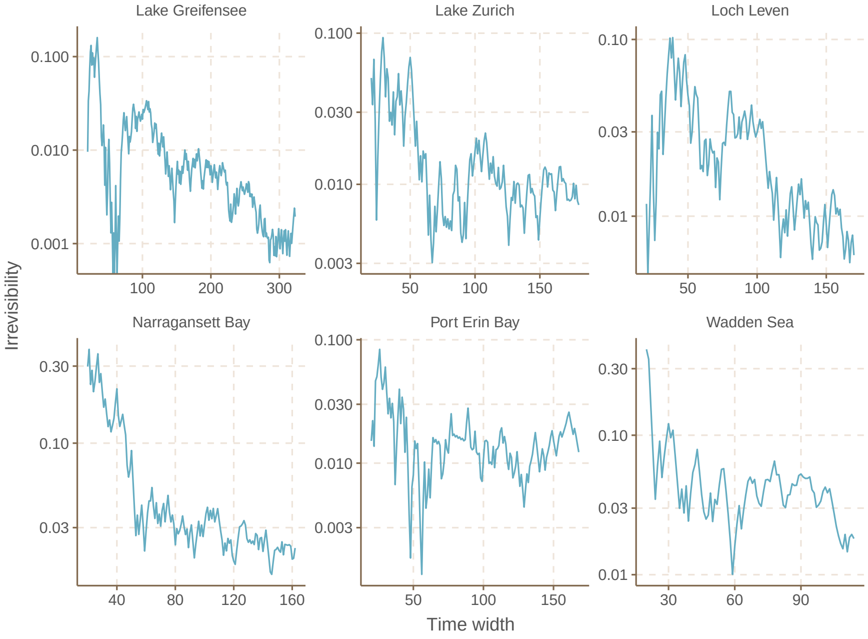


Figure S36: Convergence analysis.

Figure 4D in the main text shows that temperature and irreversibility are negatively related in these aquatic systems. Figure S37 further validates the robustness of the conclusion by varying the length of the sliding window.

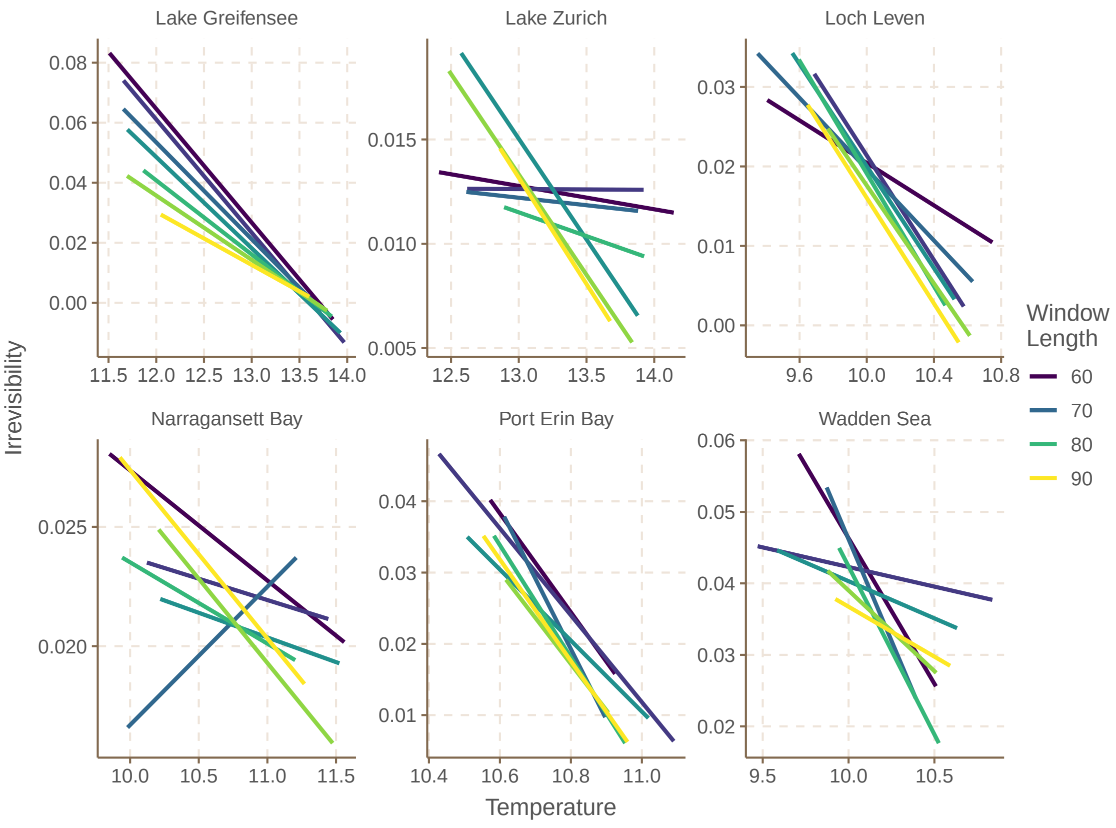


Figure S37: Generality of the negative association between mean temperature and irreversibility using varying window lengths.

### L Nonreciprocity and irreversibility

Nonreciprocity is another characteristic of nonequilibrium systems and is intrinsically linked to irreversibility. From a physical standpoint, nonreciprocal interactions among different degrees of freedom within a system indicate that these interactions cannot be derived from an energy function. Conversely, ensuring a time-reversal-invariant steady state guarantees that the system's dynamics progress downhill in a configuration space described by an energy-like function. Therefore, nonreciprocity inherently implies the break of time-reversal symmetry.

### M Table of symbols

| | |
|---|---|
| $I$ | Ecological irreversibility |
| $\mathcal{F}$ | Forward probability |
| $\mathcal{R}$ | Reverse probability |
| $\omega$ | Abundance transitions |
| $\Omega$ | All possible abundance transitions |
| $T_{\mathrm{obs}}$ | Total observation time |
| $\{x(t)\}_0^{T_{\mathrm{obs}}}$ | Time series trajectory |
| $N$, $p$ | Prey and Predator |
| $\alpha$, $\beta$, $\delta$, $\gamma$ | Coefficients to describe Lotka-Volterra model |
| $N_i$ | Abundance of species $i$ |
| $a_{ij}$ | Competition coefficient between species $i$ and $j$ |
| D | Dispersal rate |
| $\eta$ | Noise strength |
| $dW$ | Wiener process |
| $Q$, $M$, $E$ | Metabolic rate, mass, and activation energy |
| $T$ | Temperature |
| $k_{\mathrm{B}}$ | Boltzmann constant |
